# Auditory Cellular Cooperativity Probed Via Spontaneous Otoacoustic Emissions

**DOI:** 10.1101/2024.08.09.607375

**Authors:** Christopher Bergevin, Rebecca Whiley, Hero Wit, Geoffrey Manley, Pim van Dijk

## Abstract

As a sound pressure detector that uses energy to boost both its sensitivity and selectivity, the inner ear is an active non-equilibrium system. The collective processes of the inner ear giving rise to this exquisite functionality remain poorly understood. One manifestation of the active ear across the animal kingdom is the presence of spontaneous otoacoustic emission (SOAE), idiosyncratic arrays of spectral peaks that can be measured using a sensitive microphone in the ear canal.1 Current SOAE models attempt to explain how multiple peaks arise, and generally assume a spatially-distributed tonotopic system. However, the nature of the generators, their coupling, and the role of noise (e.g., Brownian motion) are hotly debated, especially given the inner ear morphological diversity across vertebrates. One means of probing these facets of emission generation is studying fluctuations in SOAE peak properties, which produce amplitude (AM) and frequency modulations (FM). These properties are likely related to the presence of noise affecting active cellular generation elements, and the coupling between generators. To better biophysically constrain models, this study characterizes the fluctuations in filtered SOAE peak waveforms, focusing on interrelations within and across peaks. A systematic approach is taken, examining three species that exhibit disparate inner ear morphologies: humans, barn owls, and green anole lizards. To varying degrees across all three groups, SOAE peaks have intra-(IrP) and interpeak (IPP) correlations indicative of interactions between generative elements. Activity from anole lizards, whose auditory sensory organ is relatively much smaller than that of humans or barn owls, showed a much higher incidence of IPP correlations. Taken together, we propose that these data are indicative of SOAE cellular generators acting cooperatively, allowing the ear to function as an optimized detector.

**Significance Statement:** The inner ear is a complex biomechanical system whose function is not well understood. To further elucidate the role of coupling in emission generation, this study systematically compares fluctuations in sound emitted spontaneously from the ear (spontaneous otoacoustic emission, SOAE) across three vertebrates. Ultimately these data serve to illustrate that the inner ear is a non-equilibrium, active system whose cellular elements work cooperatively. A clearer understanding of SOAE generation and how it manifests across the animal kingdom will significantly advance our understanding of both normal and impaired auditory function.

## 1 Introduction

### 1.1 The Active Ear and SOAE Activity

Most researchers recognize the ear to be an “active system”. That is, metabolic energy is used to improve the ear’s sensitivity and (frequency) selectivity to external sound (e.g., Recio-Spinoso and Oghalai, 2017; Altòe and Shera, 2020, see also Robles and Ruggero 2001; Hudspeth 2008; Bialek 2012 for additional considerations). This cellular-based biomechanical power amplification is generally considered to be a systems-level phenomenon, where elements collectively use energy to influence responses. Thereby, in a broader sense, the inner ear can be regarded as an example of “active matter” (Ramaswamy, 2010). An oft-cited hallmark of the active ear is the generation of spontaneous otoacoustic emission (SOAE), sound emitted by the (healthy) ear and measurable non-invasively using a sensitive microphone inserted in the ear canal (Kemp, 1979). Emission typically manifests as an array of narrow spectral peaks that are unique to a given ear and have a close relationship to perception (e.g., threshold microstructure, see Long and Tubis, 1988). Many SOAE peaks exhibit statistical properties consistent with self-sustained oscillations rather than (passively) filtered noise, thus providing evidence of the ear being a non-equilibrium, active system (Bialek and Wit, 1984; van Dijk et al., 1996; Shera, 2003; Bergevin et al., 2015).

There is a broad consensus that SOAE is primarily the result of mechanical work being done by the hair cells, which act as the primary site for mechano-electro-transduction (MET). A key impediment to our understanding of the underlying biomechanics of emission generation is uncertainty of how individual generative elements are effectively coupled together, so as to work cooperatively (Zweig, 2003). We consider cellular cooperativity as the collective effects of an active medium leading to improved function (e.g., the ear’s overall sensitivity and selectivity emerges from hair cells using energy together).2 The role of coupling in emission generation is poorly understood, especially in the face of SOAE being relatively universal across the animal kingdom. That is, seemingly disparate morphological differences at the level of the inner ear give rise to ubiquitous SOAE characteristics (Fig.1; see also Manley, 2000). Given that the biomechanics of SOAE generation are thought to be closely associated with cochlear morphology, a systematic comparative approach to identify empirical similarities and differences is desirable to elucidate core principles at work in all ears. Furthermore, this approach can yield a greater understanding of what features are necessary for a model of emission generation to demonstrably capture the underlying biomechanics of the inner ear.

**Figure 1:**
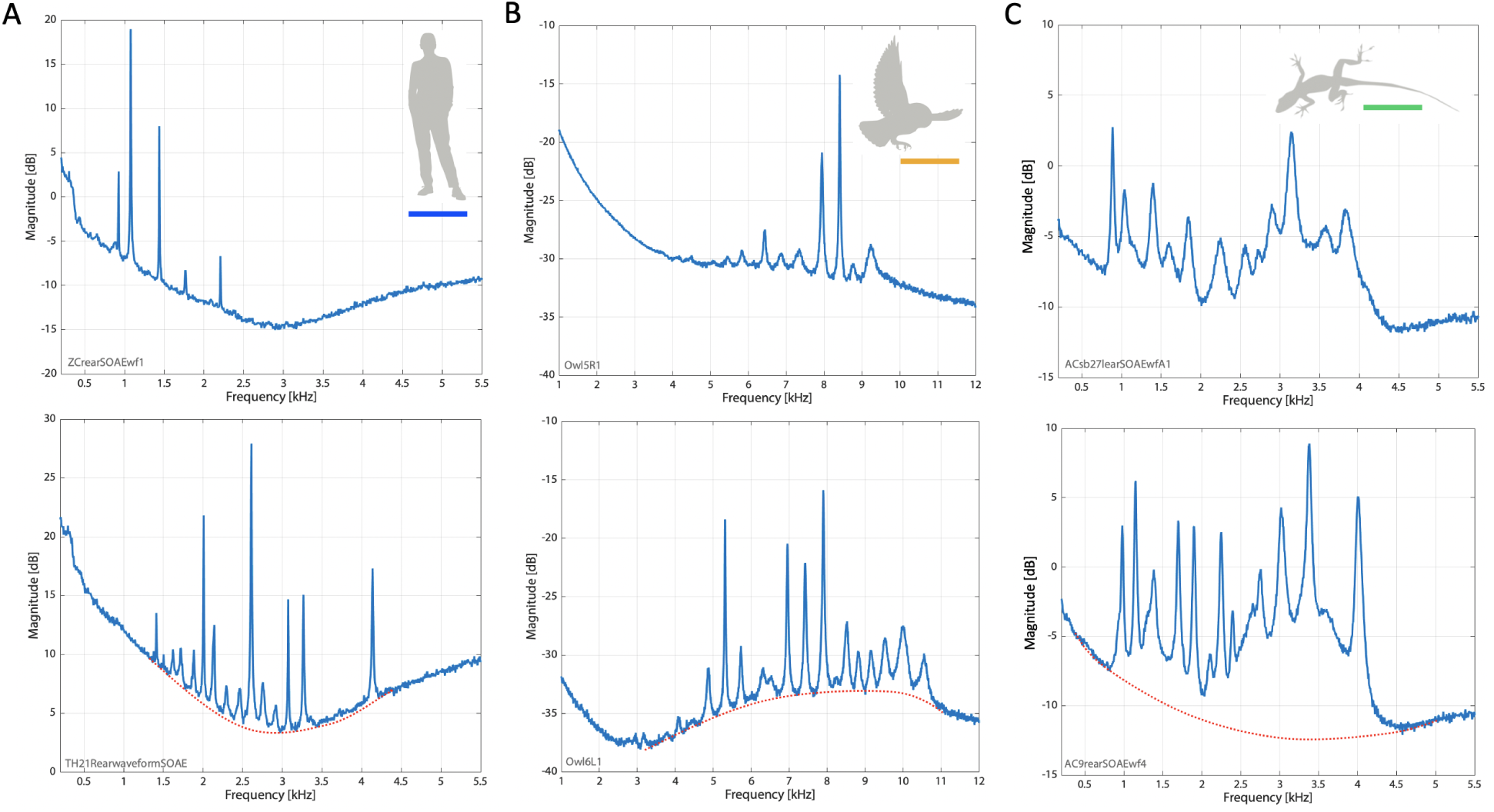
Representative SOAE spectra from humans (**A**), barn owls (**B**), and green anole lizards (**C**). Each species has data from two different individuals. The top row shows spectra from an ear exhibiting moderate SOAE activity, while the bottom row exhibits relatively robust activity. A visual approximation of the microphone noise floor, expected to be roughly similar across experiments, is shown for the bottom row (red dashed lines). For an approximate absolute reference, the microphone noise floor when coupled to a tube is typically around −20 dB SPL at 3 kHz. Ordinate units are in dB, so while there is an arbitrary vertical offset in decibels, the range is the same. See the SI (including Fig.S1) for further rationale. Short colored horizontal lines in the inset refer to the colors used in subsequent figures.

### 1.2 Modeling SOAE Generation

Historically, individual SOAE peaks were well described by first order “single-source” models, such as a van der Pol oscillator (e.g., Johannesma, 1980). Here, an implicit foundational assumption is that an individual element somehow behaves as a self-sustained oscillator (Strogatz and Stewart, 1993). Subsequent studies (e.g., O Maoileidigh et al., 2012) have expanded upon single-source models to provide possible biophysical explanations of the source of active power. However, these models are ultimately too simple to describe the unique array of SOAE peaks that emerge from a given ear, and more comprehensive approaches are needed. Shera (2022) summarized the current state of SOAE modeling and proposed two broad, but fundamentally different, categories: 1) wave-based reflection frameworks, where an otherwise stable system gives rise to SOAE somewhat analogously to a laser cavity (e.g., Shera, 2003); and 2) limit-cycle oscillator arrays that are somehow coupled to form frequency clusters, or “plateaus”, that give rise to SOAE peaks (Vilfan and Duke, 2008). Presumably this dichotomy is too simple, as they are not a priori mutually exclusive; waves may naturally arise in a limit-cycle oscillator array given coupling in a spatial-temporal system (e.g., Thipmaungprom et al., 2021), and a wave-based reflection framework can produce plateaus (Shera, 2015). Regardless of whether such a delineation clarifies the biophysical origins of emissions, the ear is a coupled system and the cellular components work cooperatively. Thus, the goal is to elucidate how this cooperativity arises.

Given the assumption that SOAE is telling us important information about how the active ear works, there is a clear need for an empirical-based thread tying together SOAE evidence to help guide modeling efforts aimed at explaining auditory function near threshold. Various characteristics of SOAE spectra, such as inter-peak frequency spacing (Shera, 2003) and the presence of “baseline activity” (Köppl and Manley, 1993; Manley et al., 1996; Manley and Gallo, 1997), have helped constrain theoretical models. One aspect that has been relatively less well characterized is the rich temporal properties exhibited by SOAE activity. For example, Burns et al. (1984) briefly reported upon an array of complex interactions between SOAE peaks that are still not well understood. While these features have been further studied (e.g., Sisto and Moleti, 1999), the temporal dynamics of SOAE is still not well understood and remain underutilized.

### 1.3 SOAE Fluctuations and a Comparative Approach

Some degree of noisiness is present in SOAE peaks, causing fluctuations in their observable properties. Specifically, a given peak exhibits time-domain variations that give rise to amplitude (AM) and frequency modulations (FM), thereby affecting the peak’s spectral shape (Köppl and Manley, 1993). A small number of studies have examined correlations between SOAE peaks (van Dijk and Wit, 1990; van Dijk et al., 1996; van Dijk and Wit, 1998a). These generally reported limited correlations for fluctuations in AM and FM within a given peak (“intrapeak”, IrP) and between AM or FM between different peaks (“interpeak”, IPP). However, a clear systematic picture has yet to emerge about the nature of what these correlations are telling us and how they arise. This is in part due to a lack of understanding as to the cause(s) of these fluctuations.

While SOAE activity is typically noted as occurring in the absence of any external sounds, the inner ear is always subject to noise.3 Such noise could drive the observed AM and FM fluctuations. When we consider SOAE as the product of cellular cooperatively, which works in the presence of noise, a critical unknown to address is the nature of the coupling between oscillatory elements. We evaluate the presence of correlations between AM and FM with the goal of providing data that can elucidate how different SOAE sources interact and clarify the distinctions (or lack thereof) between different modelling frameworks. By examining several animal groups, this study systematically characterizes IrP and IPP correlations, including how such relate to general statistical properties of individual peaks. We compare SOAE properties from three terrestrial vertebrates: humans, barn owls (*Tyto alba*), and green anole lizards (*Anolis carolinensis*). This allows us to capitalize off of the wide degree of morphological variation across vertebrate classes for properties thought to play key roles in the underlying biomechanics. Two specific considerations relevant to the three groups of this study warrant mention. First is the size of the inner ear. The human cochlea contains roughly 15000 hair cells spread over about 35 mm of length, whereas the barn owl has 16000 hair cells over 12 mm. The inner ear of the green anole contains only about 150-180 hair cell over a papillar length of 0.5 mm. Second is the presence and structure of the tectorial membrane (TM). The TM is a gelatinous, charged matrix that sits atop the stereovillar bundles of the hair cells, near the mechanically-gated transduction channels. It is considered crucial for longitudinal coupling in the mammalian cochlea (e.g., Ghaffari et al., 2007; Cheatham, 2021). In birds, the TM takes on a more massive form. In lizards, it exhibits a wide variety of shapes across classes, and is altogether absent over the vast majority (*>* 90%) of the green anole’s hair cells (Miller, 1981). Despite morphological variations, the simplest assumption is that SOAE generation mechanisms are fundamentally similar across species. We would expect that the generative elements would be affected by the same types of noise, such that AM and FM fluctuations would be similar across species, and the coupling of these elements would be preserved. Thus, one might predict that fluctuation correlations would generally be congruent across species.

## 2 Methods

### 2.1 Obtaining SOAE Waveforms

Data reported here were obtained from humans, barn owls, and green anole lizards (hereafter, anole lizards) using methods consistent with previous reports (e.g., Bergevin et al., 2015; McKetton et al., 2018; Engler et al., 2020). Human (*n* = 8) and anole lizard (*n* = 8) data were collected in Canada, while barn owl (*n* = 8) data were collected in Germany (Bergevin et al., 2015). All data were collected in an acoustic isolation booth, using an Etymotic ER-10C probe that was nonsurgically coupled to the meatus. Human subjects were awake and asked to sit motionless. Both non-human species were lightly anesthetized (owls with ketamine at 10 mg/kg and xylazine 3 mg/kg; lizards with sodium pentobarbital at 36 mg/kg). Anesthesia was maintained as necessary and animals recovered after the experiments. Body temperature was stabilized prior to recording in owls (39*^◦^*) and lizards (∼23*^◦^*) via a heating blanket. For some lizards, the heating blanket was left off (ambient temperature was usually ∼21*^◦^*). The sample rate for data acquisition was 44.1 kHz for humans and lizards, and 48 kHz for owls. Recorded waveforms were typically 120 s long (two human subjects had 60-s waveforms). Attempts were made to ensure that the obtained SOAE waveform was suitably “artifact-free” (e.g., no presence of respiratory or cardiac modulations), though not all SOAE waveforms were strictly free of artifacts (see Fig.S2). All work was approved by the York University institutional committees (protocol 2012-19) and the authorities of Lower Saxony, Germany (Niederschsisches Landesamt fr Verbraucherschutz und Lebensmittelsicherheit), permit number AZ 33.9-42502-04-13/1182. All recorded time waveforms, as well as analysis codes and filtered waveforms, are freely accessible at (*placeholder Dropbox link*) https://bit.ly/43yF0oq.

### 2.2 Averaged Spectra & Lorentzian Fitting of SOAE Peaks

Let *x*(*t*) represent the entire (“long”) waveform collected for a given ear at time *t*. The averaged spectra shown in Fig.1 were obtained by parsing *x*(*t*) into shorter consecutive segments *s* that were 186 ms long (8192 samples). The magnitude of the fast Fourier transform (FFT) of *x^s^*(*t*) was averaged for 300 segments. Thus, the spectra shown in Fig.1 were acquired over approximately one minute. For this choice of parameters, the frequency bin width was 5.38 Hz.4 To extract objective measures of peak features, we fit each peak with a Lorentzian function *L*(*f*) of the general form

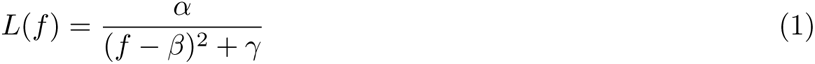

where *f* is the frequency and *β* is the peak’s characteristic frequency (CF). The function was fit to the amplitude values in dB, with a linear frequency scale. The fit was “local” in that no more than 50 frequency bins about the CF were used, and was computed via nonlinear regression using Matlab’s fminsearch.m function. An example of the fitting for all three groups is shown in Fig.S3 and further details can be found in the SI. From the fit, the peak “height” (not area) was determined as the difference between *L*(*f*) computed at the peak CF and a frequency far away from CF (e.g., 10 kHz). Peak “width” was calculated as the Full-Width Half-Maximum (FWHM) from *L*(*f*) as 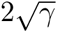.

### 2.3 Filtering of Individual SOAE peaks

Individual SOAE peaks were analyzed by computing the FFT of *x*(*t*) and applying a recursive exponential narrowband filter (see green curve in Fig.S4A, leading to the red curve by multiplication; Shera and Zweig, 1993; Shera, 2003). Filter center frequency was manually chosen to align with the frequency of maximum amplitude (i.e., the top of the peak), while the bandwidth (BW) was chosen to capture the entire width of the SOAE peak, designated as where horizontal flanks emerged atop the noise floor. Filters of adjacent peaks did not overlap. The filtered signal was inverse transformed back to the time domain (e.g., blue trace in Fig.S4B), producing a long filtered waveform for a given peak (typically being 95 s long) which was designated *x_p_*(*t*).

We used the analytic signal representation to determine amplitude (AM) and frequency modulation (FM). Let *a_p_*(*t*) represent the analytic signal of peak *x_p_*(*t*), as computed via a Hilbert transform. We obtained the AM waveform for a given peak from the magnitude of the analytic signal, commonly referred to as the envelope (red traces in Fig.S4B&D). Thus, we defined the AM of a peak as: ENV_p_(t) a_p_(t). We obtained the FM waveform by estimating the instantaneous frequency (IF) as the temporal gradient of the unwrapped phase of the analytic signal (blue trace in Fig.S4D; Boashash, 1992). Thus, we define the FM of a peak as: 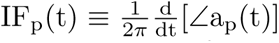. The ratio between the average IF and the filter BW was found to be close to unity within ±0.01% for all peaks, indicative that the filter center frequency serves as a suitable proxy measure of peak CF (e.g., Fig.2A), although our we ultimately used Lorentzian fits to define CF. See Table 1 for an overview of acronyms used in these analyses.

**Figure 2:**
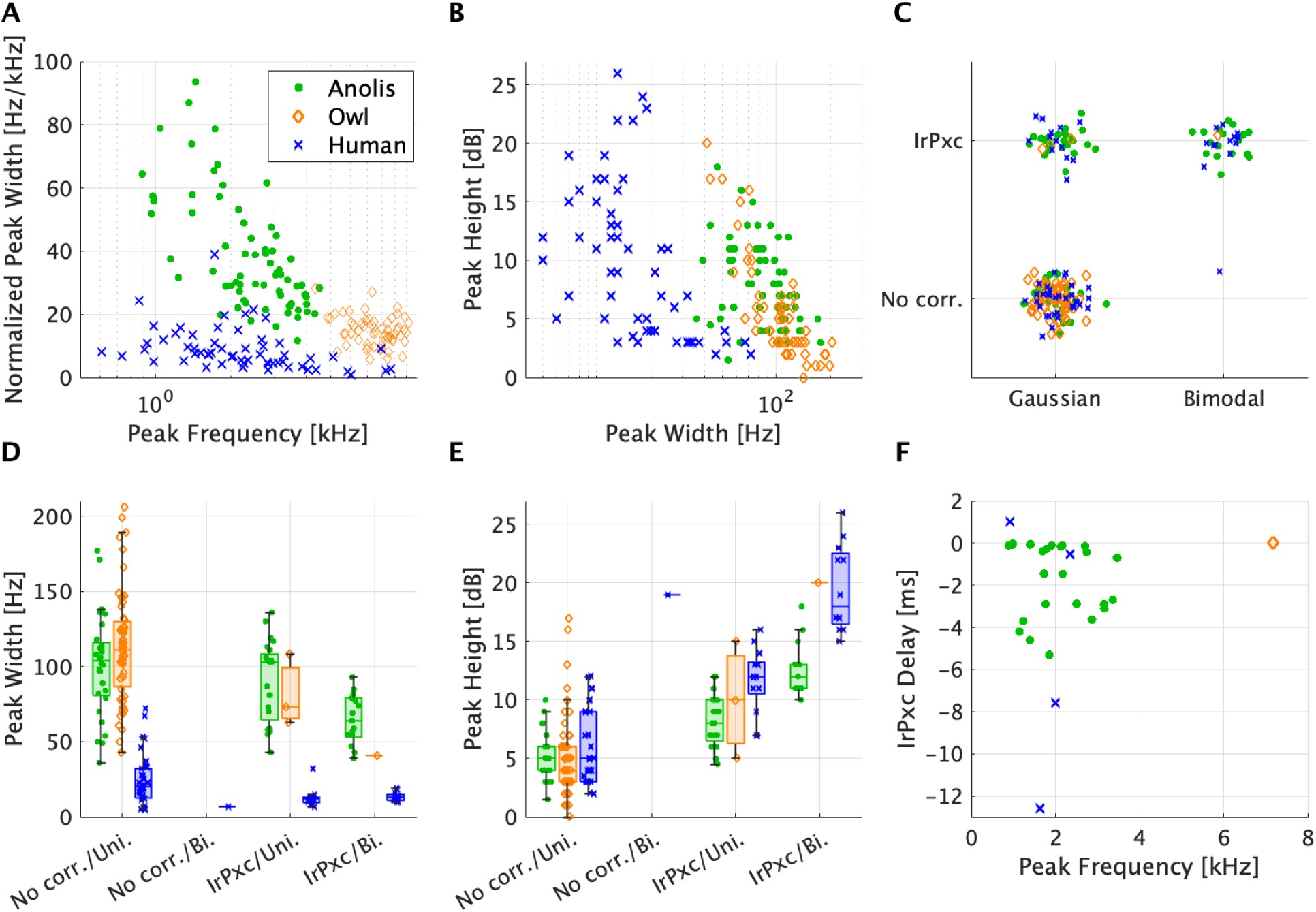
General intrapeak (IrP) features of filtered SOAE peaks from anole lizards (green circles), barn owls (orange diamonds), and humans (blue crosses). **A** – Normalized SOAE peak width (Hz) via the Lorentzian fit full-width half-maximum versus peak center frequency (kHz). Widths were normalized by dividing by center frequency (e.g., a value of 10 at 3 kHz corresponds to a 30 Hz width). **B** – SOAE peak height (in dB) versus peak width as derived from the Lorentzian fits. **C** – Comparison amongst classifiers for whether a given SOAE peak’s amplitude distribution was unimodal or bimodal (as per Hartigans’ dip statistic, *p <* 0.05) relative to whether an IrPxc was observed (|IrPxc*| >* 0.03). A small amount of vertical and horizontal jitter was added to improve visualization. **D** – SOAE peak width relative to the presence of an IrPxc and the filtered peak’s amplitude distribution, creating four categories for each species. Each box represents the interquartile range with a central mark at the median. Values beyond the boundaries of the whiskers were considered outliers. Note that no peaks from anoles nor barn owls were classified as having an IrPxc and a unimodal distribution. Only one peak from humans met this criteria, hence the lines without whiskers. There was also only one SOAE peak classified as having an IrPxc and a bimodal distribution in barn owls. **E** – Same as **D**, but for SOAE peak height across species relative to IrPxc presence and peak amplitude distribution classification. Note that in panels **D**–**E**, horizontal jitter was added to improve visualization. **F** – For SOAE peaks with an IrPxc that was localized in time (i.e., a singular peak in the cross correlation), the associated delay is plotted as a function of the SOAE peak’s center frequency.

**Table 1:**
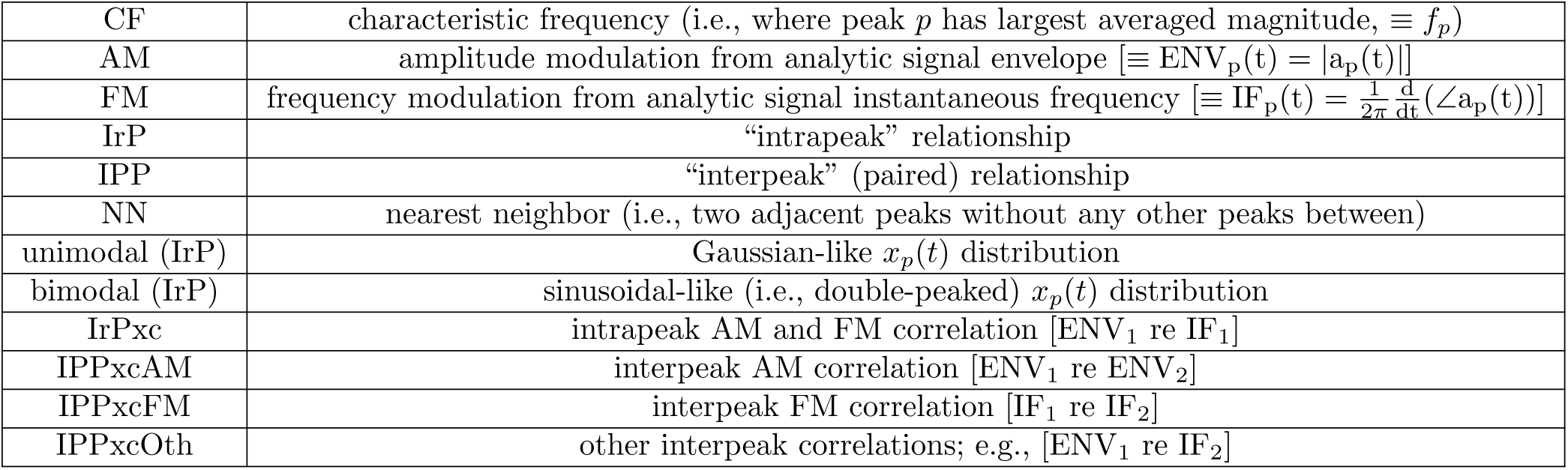
Acronyms used to describe various peak characteristics and analyses.

### 2.4 SOAE Intrapeak (IrP) and Interpeak (IPP) Analyses

Two basic types of filtered peak analyses were performed. Intrapeak (IrP) analyses concern the properties unique to a particular SOAE peak, while interpeak (IPP) concern the properties between a given pair of peaks. Peak pairs without other peaks between them were classified as nearest-neighbors (NNs; e.g., 0.93 and 1.1 kHz peaks in top left figure of in Fig.1).

For IrP analyses, we created a histogram of all values of *x_p_*(*t*) to assess whether an SOAE peak’s amplitude distribution was unimodal or bimodal (e.g., Fig.S4C). Such measures are commonly interpreted as whether a peak is indicative of filtered noise versus a self-sustained oscillation, respectively (Bialek and Wit, 1984; Shera, 2003). We classified distributions as unimodal or bimodal via Hartigans’ dip statistic (Hartigan and Hartigan, 1985), as previously described by Salvi et al. (2015), with a p-value of 0.05 across 5000 bootstraps.

This method provided a conservative estimate of distribution modality.5 We evaluated how distribution modality related to SOAE peak properties as determined by Lorentzian fitting. The second IrP analysis concerned whether a correlation was present between AM and FM traces for that peak (IrPxc). Correlation methodology is described in Section 2.5.

We performed two types of analyses to evaluate IPP relationships. First, we assessed properties related to frequency spacing between peaks (e.g., *f*_2_ *− f*_1_), including how these related to peak CF. Second, we considered potential correlations in AM or FM between peaks. Correlative behaviour between AM for two different peaks was abbreviated as IPPxcAM, whereas IPPxcFM designated correlations in FM. The analyses described here were not applied to all SOAE peaks identified. SOAE peaks identified for IPP spacing analysis were only included if the amplitude at CF was at least 1 dB above the noise floor.

### 2.5 Correlation Methodology

The potential correlations (denoted by *\**) between two isolated waveforms (e.g., AM of peak 1 as ENV_p1_(t) and AM of peak 2 as ENV_p2_(t)) were assessed to ascertain relationships between fluctuations and how in localized time such relationships are when present. We note that a key underlying assumption for standard correlation techniques is stationarity (Kantz and Schreiber, 2004). That is, whether or not the signals exhibit stability in their statistical properties over time. This is addressed further in Sec.S1.5 and the Results. While we considered various methods for correlation analyses, our primary approach was carried out in the time domain and is described here. See Sec.S1.3 for a discussion of alternate methods and cross-validations between methods.

We subtracted out the mean value of the signal and performed the correlation analyses on segments *s* of the (real-valued) residual fluctuations *φ*. For example, consider a short AM segment from a given peak 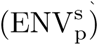 comprised of length, 𝓁, samples. Then 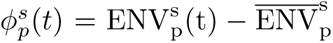, where 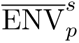 is the mean value of the envelope over the time interval spanned by the segment. Each segment was 93 ms long for lizards and owls, and 372 ms for humans due to their slower fluctuation rates (see Fig.S5), related to the generally narrower filter widths used for their peaks. For a 95 s waveform, 1022 segments were averaged for lizards and owls, and 255 segments for humans. Averaging shorter segments reduced the chance of spurious correlations occurring due to artifacts (e.g., coughing, cardiac pulsing; see also Fig.S11).

The time-domain cross-covariance between the residual fluctuations for two peaks 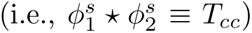 was computed using Matlab’s *xcorr.m* function as

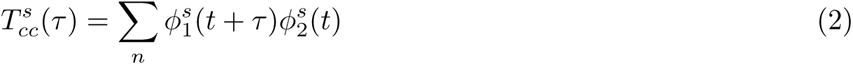

where *n* represents the total number of overlapping samples (*∼* 2*𝓁 −* 1) comprising the segments and *τ* is the “lag”. *τ* was defined such that negative values meant that *peak*_2_ was delayed relative to *peak*_1_ (i.e., temporally shifted to the right along the positive time-axis). We normalized 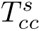 using the autocorrelations at zero lag (computed as 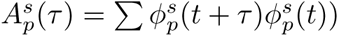 to obtain the cross correlation 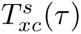 as

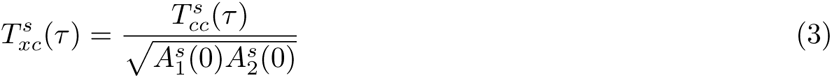

We averaged 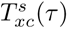 across segments to obtain an averaged, normalized cross-correlation *T_xc_*(*τ*), where *T_xc_* ∈ [−1, 1]. Negative values indicated that when one fluctuation showed an increase, the other a decrease, whereas positive values indicate both varied in a similar fashion. To cross-validate, additional correlation methods (including a spectral domain method) were also computed and were described further in the Supporting Information (Sec.S1.3).

Initial observations indicated that, when present, correlations were relatively weak (i.e., typically *T_xc_ <* 0.1), necessitating a classification metric to determine whether a correlation was present. To create a quantitative benchmark, we effectively bootstrapped the correlation to obtain *T_b_* as 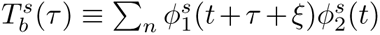, where *ξ* is a randomized indexing offset for each *τ* value to scramble the correlation reference times. We computed an averaged *T_b_*(*τ*) as done for *T_xc_*(*τ*) in Eq.3. If a fluctuation correlation was present, this had the effect of averaging it out to create control measure *T_b_*. Such a measure provides a useful visual indication of correlation presence, as shown in Fig.S6B&C (there, *T_xc_* is blue and *T_b_* is red). Second, we created an objective determination for the presence of a correlation by examining whether the maximum value of *T_xc_*(*τ*) exceeded a threshold value derived from *T_b_*. Based upon the typical range of *T_b_* values, a conservative threshold of 0.03 was typically employed (see black dashed horizontal lines in Fig.S6B&C). Rationale and justification for the choice of this value is covered in Sec.S1.3 and Fig.S8. Ultimately, this process provided a binary measure as to whether the two waveforms were correlated.

If a correlation *T_xc_* was deemed present based on the threshold criterion, it was categorized in several respects. First, we ascertained if *T_xc_*(*τ*) was singly-peaked (i.e., the cross correlation had a clear maximum or minimum localized to a particular instant; e.g., Fig.S6C). Otherwise, it was deemed indeterminate (e.g., Fig.S6B; see Fig.S13) with regard to the subsequent metrics. Second, singly-peaked *T_xc_* were classified as exhibiting positive or negative correlations based on the sign of the peak. Third, delays were determined to be positive, negative, or zero based on *τ* at the peak of *T_xc_*. A negative delay meant *peak*_2_ had to be shifted back in time to match *peak*_1_ (i.e., *peak*_2_ lagged behind *peak*_1_). We used the convention that for IrP correlations, *peak*_1_ was the AM and *peak*_2_ was the FM. For IPP, *peak*_1_ was always the peak lower in frequency relative to *peak*_2_. Taken together, these considerations form the criteria for the classifications in Table 3.

## 3 Results

### 3.1 Validation of Correlation Metholdogy

Signal stationarity is a key consideration to address, both with regard to understanding the generation sources and the primary signal processing techniques employed here, to ascertain the consistency of the statistical properties of the underlying signals across time (Hammond and White, 1996; Kantz and Schreiber, 2004). We attempted to account for potential variability in our experimental paradigm by providing time for the subject to adjust to the quiet environment in the booth before measurements (*>* 15 minutes), recording SOAE signals over a relatively short period (typically 120 s), and using waveforms without acoustical or electrical artifacts. SOAE patterns were reasonably stable across recordings when using 46 ms windows to compute a short-time Fourier transform to observe gross temporal variations (Fig.S2). When present, artifacts could be visibly discerned in the entire SOAE waveform, or identified as brief excursions in phase-plane plots of the ENV_p_ or IF_p_ signals (see Fig.S10). Analytic signal distributions likewise supported spectral stability (see Supplementary Materials Sec.S1.4); if an emission was turning on/off, we might expect a superposition of caldera and molehill distributions, which was not generally observed (e.g., Fig.S4C; see Fig.S14 for an usual case of superimposed distributions in a human subject).

We examined the autocorrelation of both the AM and FM waveforms for a given filtered SOAE peak. In general, these responses were well localized in time (e.g., Fig.S9), being strongly singly-peaked at zero lag as might be expected for a stationary noisy signal. The AM autocorrelation curves appeared Gaussian-like (i.e., continuously differentiable with zero gradient at zero lag), whereas the FM autocorrelations commonly had a singularity at zero lag (i.e., like two negative exponentials pasted together; see Fig.S9). The autocorrelations could exhibit some degree of sinc-like “ringing” (Fig.S9), which was typically small for lizards and owls but could be more substantial in humans (see peaks along the diagonal in Fig.4 denoted with !). Such may arise from the narrowband filtering of the SOAE peak.6 While there were some deviations from stationarity that likely warrant further study, especially for humans, those aspects will not be considered further here and the rest of the analyses detailed in the Results focus on methodologies that assume SOAE stationarity.

As noted in the Methods, we computed cross-correlations using both time-domain (*T_xc_*) and spectral-domain (*S_xc_*) approaches. The results reported here are exclusively from the time domain, though we typically found excellent agreement between correlation patterns for both methods. Additionally, we examined how the averaged cross-correlations for shorter time segments (i.e., *T_xc_*) compared to the response where the entire (typically 95 second) waveform was used (*T_xc_L*, where L indicates a “long” waveform used). In some cases, *T_xc_L* had a strong correlative peak but *T_xc_* did not. This typically occurred when the entire SOAE waveform contained a large broadband artifact that could allow for sufficient overlap to produce a correlation in *T_xc_L* that would be otherwise averaged out in *T_xc_*. Thus, we have no reason to expect that correlations were not detected due to the choice of shorter time segments for calculations.

### 3.2 Intrapeak Properties & Correlations

Unique spectra of SOAE peaks were apparent in all subjects, a subset of which are illustrated in Fig.1. The details of the quantity of SOAE peaks observed each species and used for spectral and correlation analyses are outlined in Table 2. Across the three species, normalized peak width decreased as peak centre frequency increased (*R* = −0.36*, p <* 0.01; Fig.2A). Generally, taller peaks had narrower widths (*R* = −0.55*, p <* 0.001; Fig.2B). As seen in Table 3 and Fig.2C, approximately one-fifth of lizard and human SOAE peaks filtered for analysis had bimodal distributions (anoles: 22.6 ± 18.8%; humans: 20.4 ± 15.9%). However, only one peak from barn owls was classified as bimodal (1.4 ± 3.9%). While a peak could be identified as unimodal as per Hartigans’ dip statistic, visual inspection of a 2-D histogram of the analytic signal could show “ringing” that was indicative of sinusoidal-like (i.e., bimodal) statistics (see right panel of Fig.S4C). The conservative nature of this classification overestimated the number of peaks that were identified as unimodal, particularly in anoles, despite having flat-top distributions that were suggestive of sinusoidal statistics obscured by noise (see left panel of Fig.S4C).

**Table 2:** Table of values from SOAE peak correlation analysis. When two values are indicated, unless noted otherwise, the first is the mean value and the second (in parentheses) is the standard deviation.

| Property | Human | Owl | *Anolis* |
| --- | --- | --- | --- |
| Total number of unique individuals considered | 8 | 8 | 8 |
| Total number of SOAE peaks considered | 73 | 94 | 87 |
| Average number of SOAE peaks/ear | 9.1 (4.6) | 11.8 (2.7) | 10.9 (2.2) |
| Number of SOAE peaks that were (Lorentzian) fit | 72 | 90 | 81 |
| Number of SOAE peaks filtered for correlation analysis | 59 | 59 | 69 |

Several examples of the AM and FM time waveforms as extracted from the filtered SOAE peaks are shown in Fig.S5. IrPxc were observed in all eight individuals for humans and lizards but were rare for owls, occurring in only two of the eight individuals. In owls and lizards, all SOAE peaks with bimodal distributions exhibited IrPxc (Fig.2C). All but one of the human SOAE peaks with bimodal distributions exhibited IrPxc. However, only a minority of SOAE peaks that exhibited IrPxc were also classified as bimodal (anoles: 41.5%; barn owls: 25.0%; humans: 48.0%; Fig.2C). We conducted three-way ANOVAs to evaluate whether peak properties were affected by amplitude distribution modality and the presence of IrPxc across species. When evaluating peak width (Fig.2D), there was a statistically significant effect of species (*F*_(2_,_171)_ = 52.3*, p <* 0.001) and a weak effect of amplitude distribution (*F*_(1_,_177)_ = 2.9*, p* = 0.09), but no effect of IrPxc presence (*F*_(1_,_177)_ = 0.3*, p* = 0.57) nor of the interactions between factors (all *p >* 0.1). Peak widths were significantly narrower in humans (19.2 *±* 14.5 Hz) than in anoles (88.4 *±* 31.2 Hz, *p <* 0.001) or barn owls (110.3 ± 37.6 Hz, *p <* 0.001), but significantly different between anoles and barn owls (*p* = 0.92). While not statistically significant, widths were generally narrower in peaks with bimodal amplitude distributions (42.2 ± 28.4 Hz) than those with unimodal distributions (79.7 ± 48.8 Hz).

All SOAE peaks from barn owls had unimodal distributions except for the tallest peak (Fig.2E). Such was in strong contrast to lizards, where peaks with much smaller amplitudes exhibited bimodal distributions. For SOAE peak height (Fig.2E), there were statistically significant effects of species (*F*_(2_,_171)_ = 15.6*, p <* 0.001), amplitude distribution (*F*_(1_,_177)_ = 25.4*, p <* 0.001), and the interaction between these factors (*F*_(2_,_177)_ = 3.05*, p* = 0.05). Across all three species, SOAE peaks with bimodal amplitude distributions were taller than those with unimodal distributions (anoles: bi., 12.5 ± 2.1 dB; uni., 6.5 ± 2.5 dB, *p* = 0.004; barn owls: bi., 20.0 dB; uni., 5.3 ± 4.0 dB, *p* = 0.016; humans: bi., 19.5 ± 3.6 dB; uni., 7.4 ± 4.1 dB, *p <* 0.001). SOAE peaks were significantly taller in humans than in anoles (*p <* 0.001), but there was no difference between humans and barn owls (*p* = 0.997). While peaks tended to be taller in barn owls than in anoles, this was not significant (*p* = 0.058). It should be noted that anole peak heights may be underestimated, since the presence of baseline activity could create elevated flanks (e.g., lizard in bottom panel C of Fig.1 has a broad hump from 2–4.2 kHz, which affects the Lorentzian fitting as shown in Fig.S3). There was no effect of IrPxc presence (*F*_(1_,_177)_ = 1.24*, p* = 0.27) nor of the interactions between IrPxc and species or amplitude distribution (both *p >* 0.10) on SOAE height.

When an IrPxc occurred for a human or an owl, it was typically not localized in time (i.e., there was not a clear singular “peak” in the cross correlation plot; see Sec.S1.5, particularly Fig.S13). For lizards, IrPxc were localized as a single peak in approximately half the cases. Across all three species, singly-peaked IrPxc were almost always negative (only one positive IrPxc was observed in humans) such that increases in AM corresponded to decreases in FM. The corresponding delays were not related to SOAE peak frequency (*R* = 0.08*, p* = 0.68). For lizards, the delays for singly-peaked IrPxc were always negative (Fig.2F), such that the FM lagged behind the AM (i.e., correlative features occurred earlier in time for AM by several milliseconds). IrPxc delays could only be determined for four peaks from humans and one peak from owls. There were numerous cases where an IrPxc was clearly present but not suitably localized in time to allow for a delay to be determined (e.g., Fig.S6B), making it difficult to characterize broad trends in the relationships between AM and FM fluctuations across species.

### 3.3 Interpeak Spacing & Correlations

We first considered “interpeak” relationships derived from the spacing between SOAE peaks in the averaged spectra (Fig.3A). The average distance between pairs of SOAE peaks (interpeak spacing) was significantly different across species (*F*_(2_,_702)_ = 22.75*, p <* 0.001). Interpeak spacing was significantly lower in anoles (1.11 ± 0.71 kHz) than in barn owls (1.81 ± 1.15 kHz, *p <* 0.001) or humans (1.70 ± 1.71 kHz, *p <* 0.001), but was not significantly different between barn owls and humans (*p* = 0.65). The distribution of interpeak ratios (computed as an SOAE peak’s CF divided by all lower peak CFs) in Fig.3B illustrates that this spacing was not the consequence of harmonic relationships between peaks. If a given peak pair was harmonically related, an integer ratio (≥ 2) would be obtained. Most ratios were less than two, with no clustering about integers, indicative that SOAE peaks were rarely (if ever) harmonic distortions of others. We computed the ratio of frequency differences amongst triplets of SOAE peaks (Fig.3C) to evaluate whether spacing was driven by peaks co-locating at cubic intermodulation distortion frequencies, which occurs when this ratio is equal to unity. While some localization about one was present for both lizards and humans, there was a broad spread suggestive that the majority of peaks were not distortions of others. However, a prominent maximum about unity was observed in owls. While decreasing the bin width demonstrated that values about 0.9-0.95 contributed this maximum, the clustering remained. This localization could be consistent with SOAE peaks being more likely to be related as intermodulation distortions in owls. Alternately, localization about one could arise by virtue of the more temporally uniform spacing between adjacent SOAE peaks in owls.

**Figure 3:**
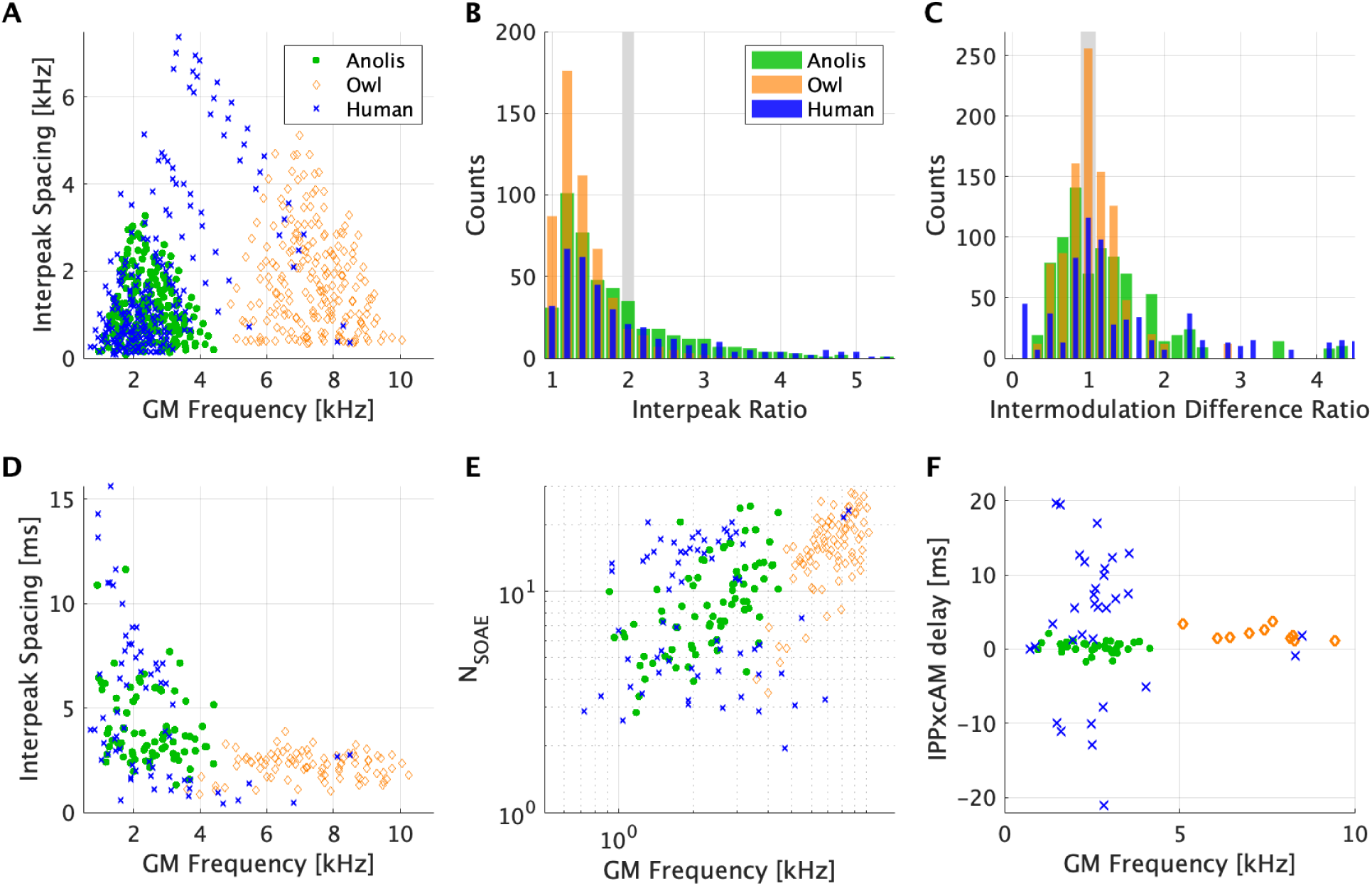
General interpeak (IPP) relationships. **A** – Interpeak frequency spacing relative to the geometric mean (GM) frequency of the pair for all SOAE peaks. **B** – Distribution of interpeak frequency ratios (i.e., CF divided by other peak frequencies below it) between all SOAE peaks from each ear. All values are greater than unity (i.e., no peaks were compared to themselves). Vertical grey line indicates a factor of two for visual reference. **C** – Distribution of intermodulation difference ratios. These were computed by considering triplets of consecutive peak frequencies (*f*_1_ *< f*_2_ *< f*_3_), counting the resulting frequency difference ratios (*f*_3_ *− f*_2_)*/*(*f*_2_ *− f*_1_). The vertical grey line, placed for visual reference, shows the ratio value that would result if *f*_1_ = 2*f*_2_ *− f*_3_ or *f*_3_ = 2*f*_2_ *− f*_1_. The total count has been divided by two to eliminate redundancies. **D** – Reciprocal of interpeak frequency spacing (hence units of ms) relative to the geometric mean (GM) frequency of the pair for adjacent peaks. **D** – Interpeak delay plotted in number of cycles (*N*_SOAE_) relative to the GM frequency of the pair for adjacent peaks. **F** – Associated delays for peak pairs with an interpeak AM cross-correlation (IPPxcAM) that was singly peaked, plotted as a function of the GM frequency of the pair. Here, a negative delay indicates that the correlated fluctuations in the lower frequency peak preceded those for the higher one.

When using interpeak spacing to estimate delays by computing the reciprocal of the frequency difference, we found that delays between adjacent peaks (i.e., NN) were significantly different across species (*F*_(2_,_227)_ = 30.83*, p <* 0.001). Barn owls had relatively uniform time delays between 1–3 ms (2.2 *±* 0.063 ms), which were significantly shorter than those in anole lizards (4.1 *±* 1.9 ms, *p <* 0.001) and humans (5.1*±* 3.7 ms, *p <* 0.001; Fig.3D). Delays in anole lizards were also significantly shorter than those in humans (*p* = 0.034). While delays were frequency-dependent in anole lizards (*R* = −0.28*, p* = 0.012) and humans (*R* = −0.45*, p <* 0.001), this was not the case in owls (*R* = 0.043*, p* = 0.6936). In Fig.3E, delays were evaluated as the number of stimulus cycles as *N*_SOAE_ (defined as the geometric mean of the pair’s CFs divided by their difference; Shera, 2003). There were still significant differences in delays between species (*F*_(2_,_227)_ = 35.6*, p <* 0.001; humans and barn owls: *p <* 0.001, humans and anoles: *p* = 0.92, barn owls and anole lizards: *p <* 0.001), but delays for owls showed greater variability and frequency-dependence in this form (*R* = 0.61205*, p <* 0.01).

Overall, a relatively small fraction of SOAE peak pairs showed interpeak correlations (Table 3). Interpeak FM correlations (IPPxcFM) were observed in six pairs in anoles (2.2% of peak pairs) and only once in humans (0.4%), but never in barn owls. IPPxcFM were only observed in SOAE peak pairs that were NNs. In anoles, IPPxcFM occurred in pairs where most SOAE peaks had amplitude distributions classified as bimodal (66.7%) and IrPxc were present (91.7%). All of the IPPxcFM in anoles had a single temporally-confined peak that was positive, such that such that increases in FM in one peak were associated with increases in the other (see example in bottom panel of Fig.4C between peaks at 1.7 and 1.9 kHz, see Fig.S6C right for correlation curve). However, their delays could be positive or negative, ranging from −0.98–2.68 ms (0.29 ± 1.40 ms). For the one IPPxcFM that was observed in a human ear, both associated SOAE peaks were bimodal but neither had an IrPxc. This IPPxcFM was not localized in time and neither a direction nor a delay could be obtained.

**Table 3:** Table of values from SOAE peak correlation analysis. When two values are indicated, unless noted otherwise, the first is the mean value and the second (in parentheses) is the standard deviation. Note that values here represent those extracted across eight different individuals for each group. Further, because not all SOAE peaks were filtered and/or all pairs analyzed for correlations, some numbers would under(over)–estimates (e.g. for anole, fraction of all possible peak pairs in a given ear showing IPPxcAM is likely lower than 17%). Several acronyms are used as follows: IrPxc *≡* intrapeak AM and FM cross-correlation, IPPxcAM *≡* interpeak AM cross-correlation, IPPxcFM *≡* interpeak FM cross-correlation, IPPxcOth *≡* cross-correlation between AM from one peak and FM from the other (and/or vice versa). Note that delay values are only included for correlations localized in time (i.e.for those singly-peaked). As note that, as described in the Methods, the fraction of peaks classified as bimodal is likely an underestimate (especially for anoles) due to limitations associated with Hartigans’ dip statistic.

| Property | Human | Owl | Anolis |
| --- | --- | --- | --- |
| **Intrapeak (IrP) Values** |  |  |  |
| Average fraction of filtered peaks/ear with bimodal distributions [%] | 20.4 (15.9) | 1.4 (3.9) | 22.6 (18.8) |
| Average fraction of peaks/ear showing IrPxc [%] | 33.9 (14.3) | 7.3 (13.7) | 58.5 (24.0) |
| Average fraction of bimodal peaks/ear showing IrPxc [%] | 88.9 (27.2) | 100.0 (0.0) | 100.0 (0.0) |
| Fraction of peaks/ear showing IrPxc and singly-peaked [%] | 3.6 (9.4) | 16.7 (23.6) | 44.9 (26.9) |
| Fraction of peaks showing (singly-peaked) IrPxc that were negatively-correlated [%] | 88.9 (27.2) | 100.0 (0.0) | 100.0 (0.0) |
| Average (signed) IrPxc delay [ms] | -4.9 (6.3) | 0.02 (0.0) | -1.7 (1.7) |
| **Interpeak (IPP) Values** |  |  |  |
| Average spacing between neighboring peaks [kHz] | 0.42 (0.48) | 0.49 (0.18) | 0.29 (0.12) |
| Fraction of analyzed peak pairs showing IPPxcAM [%] | 24.3 | 5.0 | 13.6 |
| Fraction of peaks showing IPPxcAM that were bimodal [%] | 25.9 | 10.0 | 29.7 |
| Fraction of peak pairs showing IPPxcAM that were NNs [%] | 39.3 | 70.0 | 91.9 |
| Fraction of NN peak pairs that show IPPxcAM [%] | 43.1 | 13.7 | 55.7 |
| Fraction of peak showing both IPPxcAM and IrPxc [%] | 48.2 | 15.0 | 67.6 |
| Fraction of peaks/ear showing IPPxcAM and singly-peaked [%] | 94.6 | 100.0 | 97.3 |
| Fraction of peaks showing (singly-peaked) IPPxcAM that were negatively-correlated [%] | 47.2 | 100.0 | 16.7 |
| Average (signed) IPPxcAM delay [ms] | 3.5 (9.7) | 2.1 (0.92) | 0.28 (0.77) |
| Average (unsigned) IPPxcAM delay [ms] | 8.4 (5.8) | 2.1 (0.92) | 0.63 (0.52) |
| Fraction of analyzed peak pairs showing IPPxcFM [%] | 0.4 | 0.0 | 2.2 |
| Peaks/ear showing IPPxcFM that were bimodal [%] | 100.0 | – | 66.7 |
| Fraction of NN peak pairs that show IPPxcFM [%] | 100.0 | 0.0 | 9.8 |
| Fraction of peak pairs showing IPPxcFM that were NNs [%] | 100.0 | – | 100.0 |
| Fraction of peak showing both IPPxcFM and IrPxc [%] | 0.0 | – | 91.7 |
| Fraction of peaks/ear showing IPPxcFM and singly-peaked [%] | 0.0 | – | 100.0 |
| Fraction of peaks showing (singly-peaked) IPPxcFM that were negatively-correlated [%] | – | – | 0.0 |
| Average (signed) IPPxcFM delay [ms] | – | – | 0.29 (1.4) |
| Average (unsigned) IPPxcFM delay [ms] | – | – | 0.98 (0.92) |

**Figure 4:**
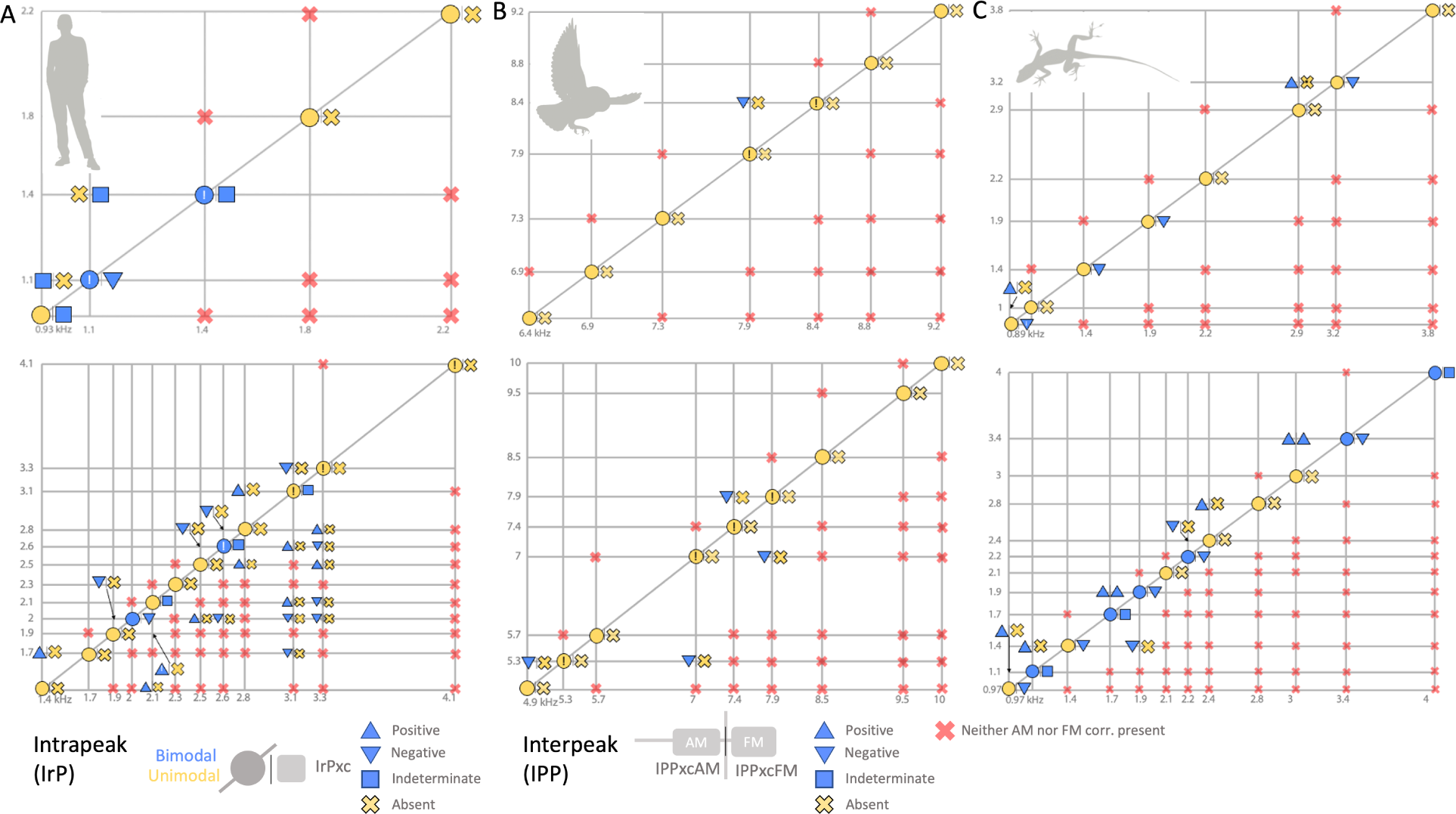
SOAE correlation maps for the same SOAE spectra (and following the same layout) as in Fig.1. Here, SOAE peak spacing is plotted logarithmically, mirrored about both the horizontal and vertical axes. Thus, if the tonotopic map is approximately exponential in nature, the line spacing would match spatial distance along the basilar membrane. Intrapeak (IrP) relations are shown along the diagonal. Nearest-neighbor interpeak (IPP) relations are in the upper half and all other peak pair relations are in the lower half. At any given intersection, a pair of symbols is present to describe the correlations, as defined visually in the legend at the bottom. The left symbol in a given pair describes the AM correlations and the right symbol describes the FM correlations. For IPP cases where neither an AM or FM correlation was detected, there is a single red x. In IrP cases where either the AM or FM waveform autocorrelations showed a significant degree of ringing, there is an ! inside the circle symbol.

While interpeak AM correlations (IPPxcAM) were more common than IPPxcFM, they still occurred in the minority of SOAE peaks for each species (anoles: 13.6%; barn owls: 5.0%; humans: 24.3%). We evaluated whether SOAE peak amplitude distribution modality or the presence of an IrPxc in a pair affected the presence of IPPxcAM across species in a three-way ANOVA. This identified significant main effects of species *F*_(2_,_695)_ = 4.72*, p* = 0.009) but not amplitude distribution (*F*_(1_,_695)_ = 0.05*, p* = 0.83), the presence of IrPxc (*F*_(1_,_695)_ = 2.27*, p* = 0.13), nor the interactions between these factors (all *p >* 0.30). IPPxcAM were significantly more common in humans (24.35%) than in anole lizards (13.55%, *p* = 0.007), but there was no difference between humans and barn owls (4.95%, *p* = 0.38) nor anole lizards and barn owls (*p* = 0.89). While many NN pairs did not show IPPxcAM, they were significantly more common between NNs in anoles (91.9%) and barn owls (70.0%), but not in humans (39.3%). We constructed “correlation maps” to combine correlation and tonotopic information visually (Fig.4). These assumed that the frequency region associated with SOAE generation (above roughly 1 kHz) has an exponential tonotopic map, consistent with auditory nerve fiber tracing studies (e.g., Turner, 1987; Manley et al., 1999). Overall, these plots indicate the relative sparse nature of interpeak correlative behavior and how such varies across groups, and also the relatively limited spatial range over which such occurs.

A visual comparison across species of both peak width and height versus frequency, relative to whether a given peak pair showed IPPxcAM or was NN, is shown in Fig.S16. A three-way ANOVA with peak width as the dependent variable identified statistically significant effects of species (*F*_(2_,_695)_ = 348.5*, p <* 0.001), IPPxcAM presence (*F*_(1_,_695)_ = 11.3*, p <* 0.001), and their interaction (*F*_(2_,_695)_ = 13.9*, p <* 0.001), but no effect of NNs (*F*_(1_,_695)_ = 0.48*, p* = 0.49) nor the other interactions (both *p >* 0.20). SOAE peaks with IPPxcAM were narrower than those without correlations in barn owls (*p <* 0.001), but not in anoles nor humans (both *p >* 0.80). We also conducted a three-way ANOVA with peak height as the dependent variable and found statistically significant effects of species (*F*_(2_,_695)_ = 10.23*, p <* 0.001), IPPxcAM presence (*F*_(1_,_695)_ = 36.5*, p <* 0.001), and their interaction (*F*_(2_,_695)_ = 9.9*, p <* 0.001), but no effect of NNs (*F*_(1_,_695)_ = 0.56*, p* = 0.46) nor the other interactions (both *p >* 0.40). SOAE peaks with IPPxcAM were significantly taller than those without correlations in barn owls (*p <* 0.001), but not in anoles (*p* = 0.90) nor humans (*p* = 0.07).

In general, human IPPxcAM peaks were wider (i.e., less temporally localized) than those in lizards or owls (Fig.S17), which is consistent with human SOAE peaks being comparatively narrower in the spectral domain (Fig.1). When IPPxcAM were deemed present, the majority of correlation waveforms (≥74%) had a single temporally-confined “peak” (e.g., Fig.S4). Other correlation waveforms exhibited more complex shapes (e.g., Fig.S12), and we could not readily infer if a correlation was positive or negative nor calculate its associated delay. For owls, all IPPxcAM were negative, whereas the majority of correlations were positive for lizards (83.3%). For humans, the fraction was roughly evenly split (52.8%). IPPxcAM delays were not frequency-dependent in any species (all *p >* 0.10), and were on the order of 0–10 ms (Fig.3F). Delays were statistically significantly different between species (*F*_(2_,_100)_ = 12.5*, p <* 0.001), being significantly longer in humans (3.5 ± 9.7 ms) than in anoles (0.28 ± 0.77 ms; *p <* 0.001) or barn owls (2.1 ± 0.92 ms; *p* = 0.006). There was no difference in delays from anoles and barn owls (*p* = 0.93). Delays tended to be positive, meaning that the SOAE peak with a higher CF led the other.

## 4 Discussion

### 4.1 General Comparison Across Groups

First we summarize the general findings from the study, drawing attention to similarities and differences across the three groups (human, barn owl, anole lizards). Each subject had a unique array of SOAE peaks. The properties of individual SOAE peaks were generally consistent across species with respect to their frequency-dependence (Fig.2). However, other peak properties were variable. Our evaluation of the effects of temporal properties (amplitude distribution modality, the presence of IrPxc or IPPxc) on SOAE peaks indicated that taxonomic class was the strongest determinant of differences in peak properties, with human SOAE peaks being consistently taller and narrower than those from barn owls and anole lizards. The presence of IrPxc had no effect on SOAE peak height or width, but IPPxcAM presence did. In particular, IPPxcAM were associated with taller, narrower peaks in barn owls, but had no relationship with peak properties in humans nor anole lizards. Across all three species, peak height was inversely related to peak width, and taller peaks were more likely to have bimodal amplitude distributions (Fig.2C&D). The relationship between amplitude distribution modality and peak height may be a consequence of higher signal-to-noise ratios in such peaks, which could explain the classification of many SOAE peak amplitude distributions as unimodal, despite features that suggested sinusoidal-like statistics in both the amplitude and analytic signal distributions. However, comparatively shorter and wider peaks from anole lizards were classified as bimodal more often than taller and narrower peaks from humans or barn owls (Fig.2C&D), indicative that signal-to-noise ratios cannot wholly explain the observed results. Rather, these variations are likely connected to morphological differences between the species examined here, further elaborated on below.

SOAE only rarely creates observable harmonic distortion (e.g., a large SOAE peak at frequency *f* does not lead to another SOAE peak at 2*f*)). However for the owl, interpeak spacing was relatively more uniform (see Fig.3C), suggestive that either intermodulation distortion was common and/or that some sort of characteristic delay was present. Despite this uniformity, there was relatively little correlative behavior between peaks (e.g., lack of clear IPPxcAM). We interpret this that peak pairs are either less likely to be a primary and associated distortion products (e.g., uniform spacing related to a standing wave effect), or some noise sources obscure such a (correlative) relation (see also van Dijk and Wit, 1998a,b). Additionally, a significant fraction of correlations deemed present were not singularly peaked. That is, their relationship could not be well localized to a particular instant (e.g., Fig.S12). For these cases, gradient-related considerations may be at play, such that the time derivative(s) of one waveform that might relate to the other (see Fig.S13). Further study is needed to explore these relationships.

There were clear differences in IPPxc across the three groups (Figs.3&4). AM correlations (IPPxcAM) were equally common for humans and lizards, but less likely for owls. IPPxcAM were always negative for owls, whereas for lizards they were mostly positive (Table 3). For humans, IPPxcAM could be in either direction. In humans, IPPxcAM delays could be tens of ms; nearly an order of magnitude longer than those observed for non-mammals (Fig.3F). While IPPxcAM delays could be positive or negative (albeit always positive in barn owls), they tended to be positive such that the SOAE peak with a higher CF led the other, which is consistent with previous observations for humans (van Dijk and Wit, 1998a).

A feature of SOAE activity that appears unique to lizards was the presence of “baseline” activity (Köppl and Manley, 1993; Manley et al., 1996; Manley and Gallo, 1997), as visually apparent in Fig.1. Baseline emissions could contribute SOAE energy in the filter in addition to the peaks, which could explain the higher incidence of correlations between smaller amplitude peaks in anole lizards. This persistent underlying broadband activity could act as a catalyst to synchronization, giving rise to both IrPxc and IPPxc. Future work should address whether correlative behavior is exclusively confined to peaks. While not systematically explored, it is possible that “inter-valley” correlative behavior could occur and could be related to the presence of the broad baseline SOAE present for anole (see Fig.S18).

The inner-ear morphological differences across the three species studied here need to be considered, as there were sufficient differences in correlative behavior to suggest SOAE generation could occur differently amongst species.7 To expand upon this point, consider how the spatial extent of the relevant portion of the tonotopic (i.e., frequency-space) map varies across the three groups. For simplicity, consider human SOAE activity as occurring over the frequency range of 0.5–8 kHz (Talmadge et al., 1993). This amounts to four octaves, which would spatially span an extent of about 15–20 mm, depending upon assumptions made about the specific form of the tonotopic map. For barn owl, SOAE occurs from about from about 2–12 kHz, or roughly 2.6 octaves (Taschenberger and Manley, 1997). In light of the underlying tonotopic fovea (Köppl et al., 1993), this amounts to a spatial distance of about 8–9 mm. Now consider the anole lizard at about 20 deg C, where SOAE frequencies are roughly 0.5–5 kHz, or 3.3 octaves. This range only spans about 0.25 mm (Manley and Gallo, 1997; Negandhi et al., 2018).8 If the longitudinal spatial extent of a hair cell can be taken as 10 *µ*m, assumed to be roughly constant across species, then clearly there are significantly fewer cells per octave for anole lizards, and thus presumably fewer cells contributing to SOAE generation. Since these cells are more closely localized together, our finding that anole lizards had a higher degree of correlative behavior is not unexpected, as interactions might be expected to decrease over longer distances. That is, spatial considerations could account for differences in the degree of SOAE peak interactions observed, including possibly explaining the presence of baseline SOAE activity.

An additional consideration that is relevant to the comparatively more “local” IPPxcAM observed in green anoles than humans or barn owls, is the organization of their hair cells. The cells are densely packed together, forming what appears as a continuum of stereovilli, akin to a phalanx (Negandhi et al., 2018). Given boundary layer considerations, this suggests that individual bundles would be predominantly viscously coupled to their nearest-neighbors. This is consistent with the observed IPPxcAM occurring most often between adjacent SOAE peaks. However, they are also embedded in the underlying papilla, which would effectively introduce a degree of “global” coupling that could affect how the cells work together (Bergevin and Shera, 2010; Bergevin, 2016; Wit and Bell, 2022). For example, the rigid papilla, which prevents the longitudinal traveling wave along the basilar membrane as present for mammals, may act as a spatial integrator to allow for a coherent response amongst contributors. Such an effect might be similar to a “a mechanical funnel” that has been proposed for localized basilar membrane regions in the mammalian cochlea Shera and Altòe (2023). Thus, this would give rise to IPPxcAM between more spatially distributed SOAE peaks.

Overall, correlations within a peak for a given ear (IrPxc) were much more common than between peaks in a given ear (IPPxcAM or IPPxcFM; Table 3). We used a threshold to make a binary classification of whether a correlation was present and did not explicitly report (normalized) cross-correlation values. When correlations were present, the values observed were typically small (~0.1 or less; see Fig.S11B) and were only attainable through averaging (or long timescales). This suggests that some source of noise obscures (or affects) how SOAE fluctuations interrelate. One interpretation is that correlations, if present, are relatively weak. Notably, SOAE signals themselves are typically small in terms of filtered energy in the “peak” relative to the underlying noise floor (see Fig.1 of van Dijk et al., 1996), which would likely affect the observed correlation strength. More IPP correlations may emerge (especially for humans) when using improved time-series analysis techniques, including both linear (Middleton, 1960) and nonlinear methods (Kantz and Schreiber, 2004). These could include modern entropy-related analyses (e.g., maximum entropy spectral estimation) (Skinner and Dunkel, 2021; Roldán et al., 2021; Sheth et al., 2021) and higher-order spectral analysis (Nikias and Mendel, 1993). Further, inclusion of techniques specifically addressing non-stationarity (Kantz and Schreiber, 2004; Hammond and White, 1996) could prove valuable.

SOAE peaks appeared stationary to a first degree, but human SOAE had greater deviations from stationarity than anole lizards or barn owls. The narrower peak widths observed in humans could underly these deviations. While not systematically explored, anecdotal observations indicated that the choice of filter bandwidth did have some effect upon the observed correlations. As long as the filter width captured the bulk of the energy about the SOAE peak, making the filter slightly narrower or wider had little effect upon the qualitative shape of a correlation (if present), causing only slight changes in the correlation height or delay. This occurred even despite clear visual changes in the fluctuations (e.g., widened filters for human peaks meant faster fluctuation waveforms that differed by visual comparison to the standard narrower filters). Several different correlation methods are shown in Fig.S7, indicating that all lead to qualitatively similar endpoints, except in rare instances. These observations suggest filter parameters do play an important role, but do not critically affect the correlative features as reported here.

### 4.2 Biomechanics Underlying Spontaneous Oscillations

Given that we interpret the data reported here as indicative that SOAE generators are acting cooperatively, we next frame the general question(s) about what is (un)known in terms of structures vibrating spontaneously. Ultimately, very little is currently known empirically about what is actually oscillating spontaneously inside the inner ear. For example, the following questions remain open:

1. What is the source of the oscillations in the inner ear that generates SOAE activity?
2. How do these oscillations arise given the spatially-distributed nature of the inner ear?
3. How are generative elements coupled together so to affect each other’s behavior?
4. What causes the (amplitude and frequency) fluctuations in SOAE activity, giving rise to a peak’s width?
5. What are the relevant sources of noise (e.g., Brownian motion, channel clatter) that affect SOAE generation?

Given that these are foundational questions with regard to the modeling assumption underlying SOAE generation, we briefly review what has been reported. Nuttall et al. (2004) directly observed spontaneous BM motion in a guinea pig that corresponded to SOAE measured in the ear canal. This fortuitous observation was made in a single animal and unfortunately has not been replicated to the best of our knowledge. Nuttall et al. (1997) found that BM (basilar membrane) motion exhibited noise that was band limited to the tonotopic measurement site and was also affected by physiological factors (e.g., decreasing with or and efferent stimulation). However, the source of the noise remains unknown; it could be caused by external sound pressure on the eardrum, internal Brownian motion of fluid, and/or something else entirely. Spontaneous vibrations of the eardrum for anole lizards have been shown to directly correlate to SOAE (Bergevin et al., 2018). Additionally, indirect measurements offer further insights on spontaneous oscillations in the cochlea (e.g., Powers et al., 1995; Cheatham et al., 2014; Quiñones et al., 2022). For example, Cheatham et al. (2014) found that genetic mutations that affect the tectorial membrane significantly affected the presence of SOAE in mice. Lastly, one influential reported measurement is the observation of spontaneous oscillations of bullfrog saccular hair bundles (e.g., Martin et al., 2003; Salvi et al., 2015), which have formed the basis of one class of SOAE models, namely the “local oscillator” group. However, it is unclear if these types of motions, which occur at tens of Hz, extend to the auditory range of several kHz or higher where fluid forces may be very different (Freeman and Weiss, 1988, 1990; Kozlov et al., 2011). Consider for example that SOAE activity has been observed up to 63 kHz in bats, a frequency region important for echolocation in that species (Kössl, 1994).

### 4.3 Connecting Models to SOAE Fluctuations

To help bridge the last two sections, we briefly describe the current state of SOAE modeling and frame how the present results can inform such moving ahead. In general, cochlear models do not address SOAE generation. Older model classes for “simpler” ears tended to be passive, one comprehensive example being a cascade of linear time-invariant filters including a static nonlinearity (Weiss et al., 1985). While it is still unclear whether these models can plausibly capture the sharp tuning and nonlinear features observed physiologically, they certainly are not capable of describing SOAE generation. Subsequent models included some sort of active contribution, though this would manifest in a wide variety of different ways (e.g., regions of negative damping, hair cell bundles acting as limit-cycle oscillators). Early SOAE models often focused on a single peak described by a single limit-cycle oscillator (e.g., van der Pol, Johannesma, 1980), including noise to create fluctuations (Bialek and Wit, 1984). These models were able to capture many basic features of isolated SOAE peaks, such as their general Lorentzian shape. Later SOAE models incorporated two or more self-sustained oscillators (e.g., Murphy et al., 1995a), leading towards several classes of models that attempt to explain multiple SOAE peaks. These models can be heuristic (e.g., Shera, 2003), computational (e.g., Vilfan and Duke, 2008; Gelfand et al., 2010; Epp et al., 2010; Bowling et al., 2019), or analytically based (e.g., Talmadge et al., 1998; Ku et al., 2008), but none have been reported as being able to reasonably capture the characteristics of SOAE fluctuations.

To frame how the current results can help constrain models of SOAE generation, we briefly expand upon the proposed modeling dichotomy (Shera, 2022) described in the introduction. The following rhetorical question serves as a launch point: Where is the cochlea/a hair cell “poised”? (Zweig, 2003). In essence, this distinction revolves around whether an individual element can self-oscillate (i.e., and thereby the spontaneous oscillation is “local”; see Hudspeth (2010)), or whether it is necessary to embed the element into a distributed system for such spontaneous oscillations to manifest (and thus it is “global”). Applied to SOAE generation, the “local” limit-cycle oscillator array formulation assumes that the various generation elements individually behave as limit-cycle oscillators. Conversely, the “global” wave-based reflection framework posits that SOAE is intrinsically an emergent systems-level response; the (inner) ear is a collective system that is wholly different than the sum of the parts. In essence, this dichotic framework ties back to arguments about the parameter regime for models.

Ultimately, we would argue that this proposed distinction is circular, as it is predicated upon the unknown answer to the question of precisely what is oscillating and what are the forces causing such.9 For example, a “system” of limit-cycle oscillators can sit in an “amplitude death” regime when coupled together (Murphy et al., 1995b; Ahn, 2013)10 Thus a limit-cycle oscillator array need not intrinsically oscillate spontaneously. Further, the distinction is a potentially confounding choice of wording: “global” versus “local” can mistakenly be interpreted as referring only to (inter-element) coupling.

Nonetheless, examples of each modeling approach have both merits and drawbacks. A standing wave-based reflection framework (Shera, 2003) has made several predictions that reasonably hold, although it has not been placed in a more comprehensive computational scheme to test how robust the model is for predicting a wider range of SOAE phenomena. A limit-cycle oscillator array (Vilfan and Duke, 2008) describes a variety of features via “frequency clustering”, but does not yet produce realistic SOAE spectra (e.g., peaks are too narrow). Further, to reiterate, it is unknown if the hair cells in non-mammalian ears do in fact act as “local” oscillators. Thus, a key central assumption has yet to be empirically validated. In summary, the distinction can be stated quite colloquially: Is the heuristic one of “SOAE generation” or “SOAE generators”?

We propose that the data reported here demonstrating the presence of both intra- and interpeak correlations are indicative of different SOAE sources interacting. Further, we propose that the cellular cooperativity that presumably leads to SOAE, works in the presence of noise (e.g., Brownian motion). Seen through this lens, the two proposed model classes appear just as different facets of the same essential contrivance: a theoretical description of a spatially-distributed system of coupled active (i.e., some sort of internal energy source) oscillatory elements. The key challenge at present is to properly describe that coupling so to elucidate how the cellular elements cooperatively work together to create self-sustained motion, giving rise to individual SOAE peaks, and further characterize the precise role of key aspects such as nonlinearity and noise. For instance, assuming some source(s) of noise that causes SOAE fluctuations, one might expect differences in correlative behavior across models: a limit-cycle oscillator array viewpoint (e.g., nearest-neighbor coupling) might expect correlations to be localized to neighboring peaks, whereas a wave-based reflection framework perspective might predict more broader correlations due to a noisy reflection boundary that is shared across generation sites (e.g., the stapes as driven by external noise affecting the eardrum).

To help guide future SOAE modeling efforts, we pose several lines of inquiry stemming from the current results. Some of these revolve around what sort of inter-element mechanical coupling should be considered, as well as what sort of role stochastic considerations play:

1. What is the dominant source of noise to which an individual generator element is subject (e.g., Brownian motion affecting displacement, channel “clatter” in the mechanically-gated ion channels, etc.)?
2. Given the relatively limited tonotopic extent of correlative behavior (i.e., predominantly confined to nearest-neighbor peaks), is the basis for energy flow diffusive or wave-based in nature (Mandelis, 2000)? More generally, how spatially localized is the noise, such that forces experienced are correlated to some extent along the tonotopic axis?
3. To what extent can stochastic processes be used to understand the effective “bandwidth” of SOAE peaks, and why are human peaks relatively so narrow?
4. Why is correlative behavior, while present, rare? Is it because (weaker) correlations may in fact be present, but some source of measurement noise serves to mask the effect? Or are there only correlations between specific generators, perhaps due to some underlying network connectivity (e.g., nearest-neighbors, Fig.4)? Are “chimera states” [e.g., Abrams and Strogatz (2004); Faber and Bozovic (2021)] a heuristic to pursue?
5. Despite the ear likely having sufficient (nonlinear) complexity, SOAE appears generally stable with regard to external stimuli (Fig.S21). This suggests it is less likely that models should exhibit chaotic dynamics (e.g., peak structure is sensitive to “initial” conditions).
6. How can SOAE interactions inform how “roughness” (Zweig and Shera, 1995; Vilfan and Duke, 2008) should be properly included so to lead to a testable predictions?

Ultimately, these questions can also be viewed as a means to help guide models in terms of their falsifiability (Zweig, 2003): a given model can be tested against the SOAE interactions as described in the current paper.

### 4.4 Role of Nonmammals in SOAE Research

Lastly, we briefly argue that non-mammalian models are particularly well suited for future SOAE experiments and modeling, as well as auditory neurophysiology, especially when placed in a broader evolutionary context (e.g., Losos, 2011). By clarifying processes in these relatively simpler ears, key principles should become apparent that constrain and inform assumptions for the more complex mammalian cochlea (Manley, 2000). With respect to barn owls, they are known to be auditory specialists with exceptional sound localization acuity and have a well-established history as a model for auditory research (e.g., Carr and Konishi, 1990; Konishi, 1973; Köppl, 1997; Taschenberger and Manley, 1997). The barn owl’s auditory epithelium is also notable in that almost half of its length is devoted to the highest octave (from 5 to 10 kHz), thus an unusually large number of hair cells will be responsive to such frequencies. It is presently unclear to what extent SOAE activity can be observed in other bird species however.

Future work in lizards could be a particularly valuable avenue to characterize how various morphological traits could be associated with fluctuation correlations and their properties. First, consider that papillar morphology varies significantly across lizard families (e.g., number of hair cells/ear, structure/presence of tectorial membrane; see Wever, 1978; Miller, 1981; Manley, 2000). This variability motivates the application of the correlation methodology employed here to study SOAE interpeak correlations for other lizard groups. Second, we further rationalize the choice of *Anolis* as lizard for the SOAE data presented here. Anoles are an iguanid species with a small basilar papilla containing relatively few auditory hair cells (≈150), whose bundles are predominantly free standing (i.e., lack an overlying tectorium). The cells are densely packed together, forming what appears as a continuum of stereovilli, akin to a phalanx (Negandhi et al., 2018). Given boundary layer considerations, this suggests that individual bundles would be predominantly viscously coupled to their nearest-neighbors. However, they are also embedded in the underlying papilla, which would effectively introduce a degree of “global” coupling that could affect how the cells work together (Bergevin and Shera, 2010; Bergevin, 2016; **?**). Given morphological differences relative to the mammalian cochlea, the very small ears of anole lizards bring into focus the important question central to this study: How does energy spatially propagate throughout the inner ear to affect collective dynamics? For example, the rigid papilla, which prevents the longitudinal traveling wave along the BM as present for mammals, may act as a spatial integrator to allow for a coherent response amongst contributors. Such an effect might be similar to a “a mechanical funnel” that has been proposed for localized BM regions in the mammalian cochlea Shera and Altòe (2023). Ultimately, it is presently unclear what role waves play in the anole lizard’s inner ear, if any.11 Lastly, lizards provide an opportunity to consider acoustic crosstalk between the two ears via the interaural canal (Roongthumskul et al., 2019; Bergevin et al., 2020). This pathway allows for mechanical coupling between the two ears, which broadens the notion of “global” across the head to the bilateral domain and how such may affect SOAE correlation analyses.

## 5 Summary

The present study took a comparative approach to systematically characterize SOAE peak fluctuations and their associated correlations. More specifically, we examined three species whose inner ears differ characteristically in their anatomy. Key differences include the size of the epithelium (from 0.5 to 35 mm in length) and the corresponding number of hair cells (150 to 15000), both of which presumably affect the coupling of hair cells. For example, whereas hair cells that respond to frequencies where peaks occur in the SOAE spectrum completely lack a tectorial covering in anole lizards, both barn owls and humans possess a thick, continuous tectorium. All three types of ears show SOAE activity as an idiosyncratic array of peaks of varying heights and widths. These peaks can show clear evidence of self-sustained sinusoidal oscillation (i.e., bimodal amplitude distributions), a hallmark feature of the active ear. However, the extent to which this characteristic was present was variable, being more common in humans and anole lizards than in barn owls. Further, SOAE peaks from all three groups exhibited fluctuations in amplitude and frequency that could show both intra-(IrP) and interpeak (IPP) correlative behavior. Overall, these correlations were rare in barn owls. IrP behavior was most common in anole lizards and moderately present in humans. Conversely, IPP correlations were most common in humans and less so in anole lizards. This latter IPP aspect, when present, was generally localized to adjacent SOAE peaks in barn owls and anole lizards. However, in humans, IPP behavior was more likely occur between more distant SOAE peaks than those that were directly adjacent.

We argue that IPP correlations could be attributed to how coupling might be affected by spatial aspects such as number of hair cells and of inter-cell spacing, with anole lizards at one end of the coupling spectrum and humans near the other end. While many questions still remain about the physiological underpinnings of SOAE generation, we interpret the data as indicative of SOAE arising by virtue of cooperative behavior, where cellular elements work together as active force generators to improve both the sensitivity and selectivity of the ear.

## 6 Acknowledgements

Supported by the Natural Sciences and Engineering Research Council of Canada (NSERC) to CB and RW. Input from Andrew Bell, Julien Meaud and Daibhid O Maoileidigh is gratefully acknowledged.

## S1 Supporting Information

### S1.1 Methods: SOAE Analysis & Peak Fitting

Figure S1 indicates how the absolute value of the SOAE spectra ordinate depends in part upon the length of the fast Fourier transform (FFT) time window used. Away from SOAE peaks, doubling the length of the window has the effect of halving the frequency bin widths, leading to a drop of one half (or 3 dB), as observed. Thus, as would be expected for a noisy signal, the absolute value of the noise floor depends upon the chosen signal analysis parameters. Further, near SOAE activity (especially close to the top of the peaks), the picture can become further complicated. For the range of window lengths shown in the right panel of Fig.S1, a 12 dB expected span is only about 7–8 dB. Such observations are generally present for human and owl SOAE spectra as well. These observations indicate the challenge associated with specifying absolute values for SOAE peak heights in units such as dB SPL.

**Supplementary Figure S1:**
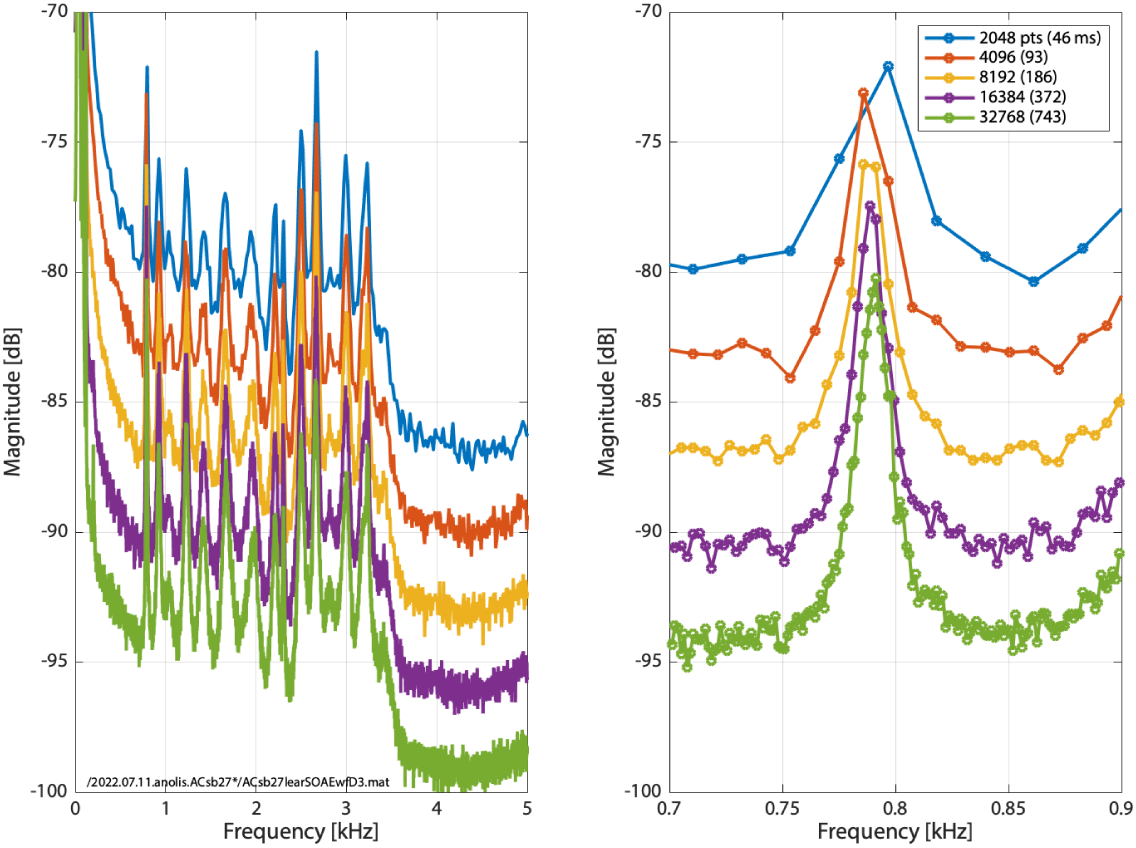
Example for an anole lizard SOAE spectrum to illustrate how response magnitudes depend upon the FFT segment length for a given sample rate. Here, the SOAE waveform was obtained at 44.1 kHz, but the window length (and the corresponding duration in ms) used for computing the FFT varied as indicated in the legend (different colours). In all cases, 150 segments were spectrally averaged to produce the shown spectra (and changing that value does not significantly affect the trend shown here). The right panel provides an inset of the left panel focussing on the SOAE peak between 0.7–0.9 kHz.

Overall stability of SOAE spectra can be observed in Fig.S2. In the top row, the SOAE peaks can be seen as horizontal bands that appear reasonably stable across the 46 ms timescale window used for the FFT. These observations provide reasonable support for the assumption of stationarity underlying the correlation analyses of the study (see also Fig.S9). In some instances, these heat maps can show artifacts present in the signals (e.g., Owl5R1 appears to show a broadband respiration artifact with frequency 70 events per minute).

Figure S3 shows examples of fitting Lorentzian functions to SOAE spectra to determine peak height and width. The fitting function was

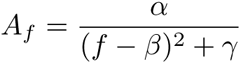

where *A_f_* is the fit amplitude, *f* is frequency, and [*α, β, γ*] are the three free parameters for the fit. Typically, only 15–25 data points on either side of the peak were used to determine the degree for which the fit was “local”. Comparisons were made between visual estimates of peak height and fit values and good agreement was found. Further, the filter bandwidth and fit FWHM were found to correlate reasonably well (i.e., filter bandwidth was a reasonable proxy measure for peak “width”). In some rare instances, when the fit was obviously unacceptable (e.g., note the owl peak in Fig.S3 just above 8 kHz), the number of points used for the fit was adjusted or the peak was otherwise altogether excluded.

**Supplementary Figure S2:**
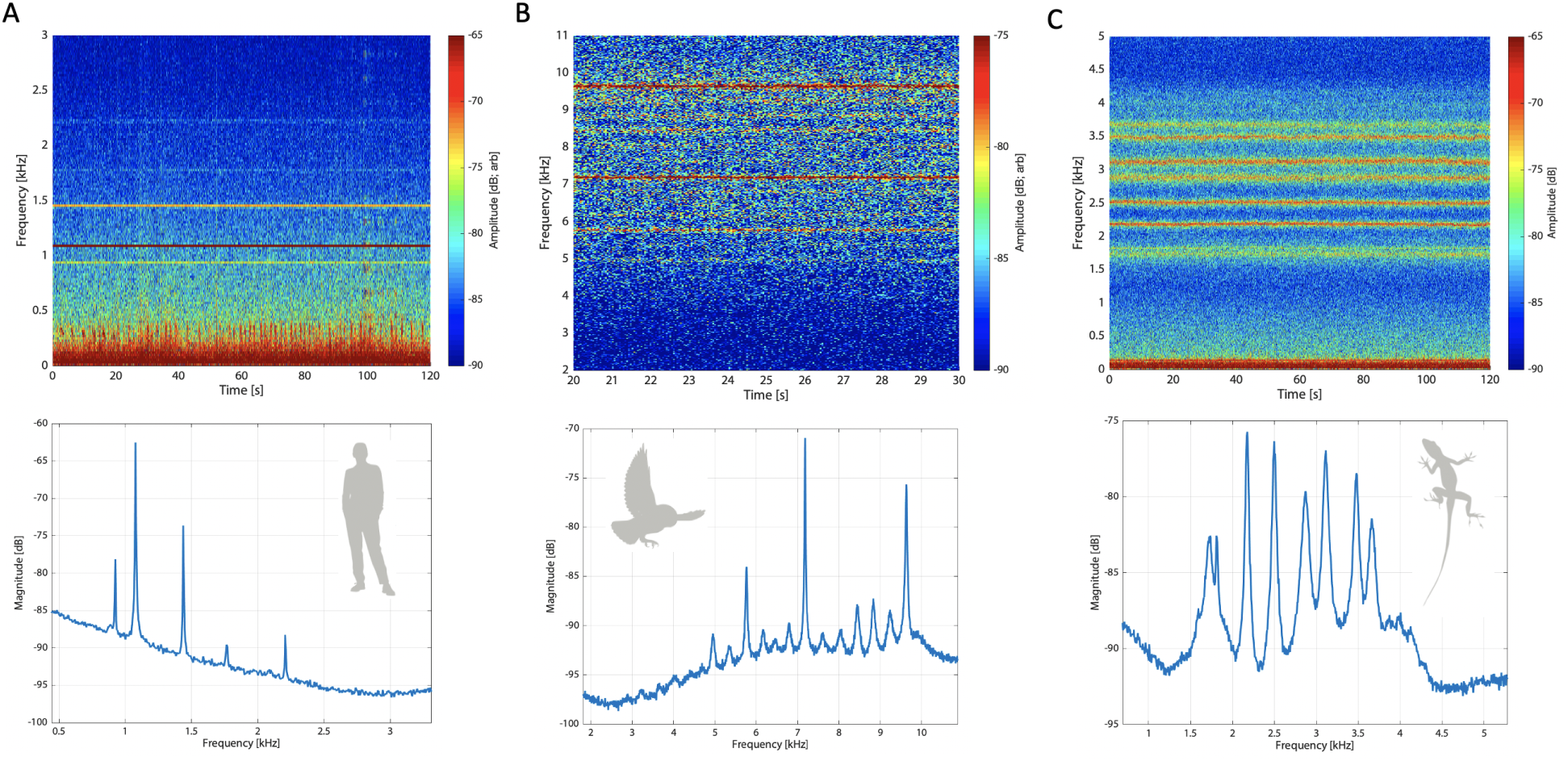
Examples of SOAE spectral variations across time by plotting as a temporal heat map (i.e., a spectrogram via a short time Fourier transform; top panels) and the corresponding SOAE spectra (bottom panels). An example from each group is shown: **A** – Human (ZCrearSOAEwf1). **B** – Owl (TAG4learSOAEwf1). **C** – Anole (ACsb24rearSOAEwfA1). A window length of 46 ms was used, with no filtering nor overlap between adjacent windows.

### S1.2 Methods: Peak Filtering & Extracting AM and FM Fluctuations

The peak filtering and cross correlation processes are illustrated in Figs.S4 and S6, respectively. Figure S4A shows a representative SOAE spectrum from anole lizard with a filtered peak at 2.178 kHz. The time waveform for this peak, and its associated envelope, are shown in panel B. The amplitude distribution of this peak is illustrated as a histogram in the left panel of C, which is classified as uni- or bimodal as per Hartigans’ dip statistic (Hartigan and Hartigan, 1985). The right panel of C illustrates the peak’s analytic signal distribution, which exhibits ringing (i.e., the count increases before decreasing when moving away from the origin at [0,0]) that is indicative of sinusoidal-like statistics. Despite the clear ringing in the analytic signal distribution and the flat-top and slight dip about zero Pa in the amplitude distribution, the peak was classified as unimodal by the dip statistic. Sinusoidal-like statistics in noisy peaks may be better captured by ringing in the analytic signal distribution than the amplitude distribution. Computing the analytic signal of the time waveform yields the AM (red solid line) and FM (blue dashed line) waveforms as shown in panel D. Additional AM and FM waveforms for representative pairs of peaks from all species can be seen in Figure S5. With these waveforms in-hand, various cross-correlations within a peak (IrPxc) and between peaks (IPPxc) could be computed as shown in Fig.S6.

### S1.3 Methods: Additional Correlative Methods

While the main document reports time-domain correlation using averaged segments (*T_xc_*(*τ*)), two additional methods were explored to cross-validate. First was the “*T_xc_L*” method, similar to *T_xc_* except that the entire (long) waveform was used rather than shorter segments, and thus there was no averaging. Second was a spectral domain method (“*S_xc_*”) that has been previously used (e.g., van Dijk and Wit, 1990, 1998a) and is described briefly here. Once two given time segments had been isolated and the mean subtracted to obtain 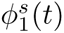 and 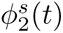, their Fourier transforms were determined via a FFT rather than computing the cross-correlation in the time domain. For *peak*_1_ this is defined as 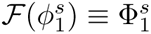 (and similarly 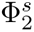 for *peak*_2_). Then we define the spectral domain cross-covariance *S_cc_*(*τ*) as

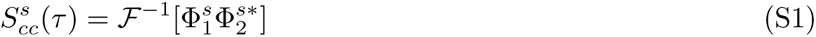

**Supplementary Figure S3:**
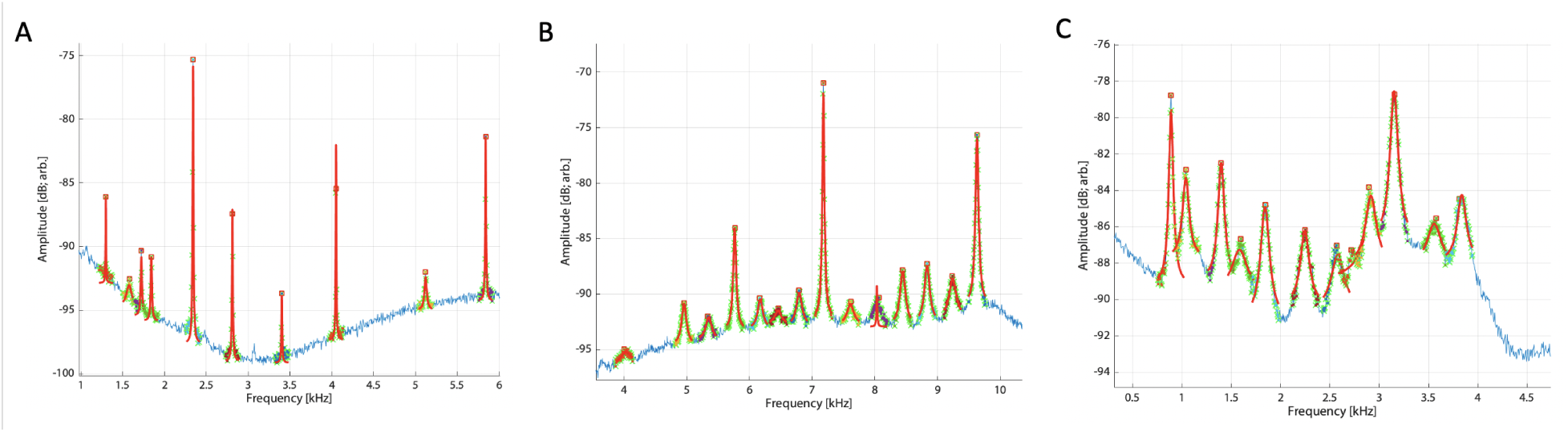
Examples SOAE peaks locally fit with Lorentzian functions via nonlinear regressions. **A** – Human (JIrearSOAEwf2). **B** – Owl (TAG4learSOAEwf1). **C** – Anole (ACsb27learSOAEwfA1). Blue trace is the SOAE spectra, green crosses indicate the local points used for a given peak’s fit, red squares represent the pre-specified peak center frequency, and the red curves are the actual fits. From the fit, the peak “height” (i.e., difference between maximum and the floor) and bandwidth (as the FWHM) could be determined. Note that not all fits were successful (e.g., peak just above 8 kHz in panel B) and thus were excluded.

where the * indicates the complex conjugate and *F^−^*^1^ the inverse Fourier transform. Thus, in the spectral domain, the correlation is simply acquired by a single (complex-valued) multiplication. We then normalized using standard deviation to obtain the spectral cross correlation 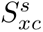 as

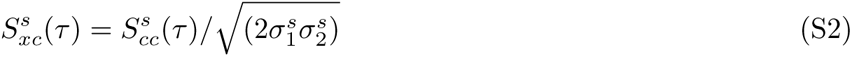

where 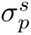 represents the standard deviation of 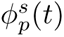 over the time segment *s*. Like the time-domain case, we then averaged across segments to obtain *S_xc_*(*τ*).

While not explored in quantitative detail, the three methods typically yielded similar results, as shown in Fig.S7. In some instances, for human IPP curves exclusively, differences were observed between *T_xc_L* curves relative to *T_xc_*, where the latter showed no IPPxcAM but the *T_xc_L* was clearly peaked. This sort of effect, only observed rarely, could linger for nearby peaks, but decreased as one moved away in frequency. In those instances, the IPP pair was considered not correlated due to the absence of *T_xc_* (with regard to Table 3). The cause for this was not precisely determined, but may be due to inclusion of an artifact that contaminated the waveform. Correlation delays could also be extracted from the spectral method using the phase gradient, as described in (van Dijk and Wit, 1998a). Preliminary observations indicated that such a method was more challenging to implement, though yielded consistent measures as the time-domain approach.

Figure S8 provides a visual justification for the choice of 0.03 as the threshold value for identifying significant correlations. When no correlation was present, the correlation values had distributions clustered about zero (panel C), whereas singly-peaked correlations showed clear asymmetric “tails” in the positive or negative direction (non-singular peaks could show tails going either way). The threshold value of 0.03 used in this study was chosen to encompass these tails, while avoiding small spurious ones from the control response.

Lastly, we also pseudo-simulated SOAE waveforms by creating noisy sinusoids with varying degrees of AM, FM, and/or additive noise. Such provided useful benchmarks to validate the computations and verify various measures (e.g., sign of delay indicating which waveform lagged the other).

### S1.4 Results: Stationarity

Signal stationarity is a key consideration to address, both with regard to understanding the generation sources and the primary signal processing techniques employed here, to ascertain the consistency of the statistical properties of the underlying signals across time (Hammond and White, 1996; Kantz and Schreiber, 2004). There are many factors that could influence SOAE stationarity, both outside the inner ear (e.g., cardiac and respiration artifacts, middle ear effects) or from within (e.g., slow adaptation inside the isolation booth, efferent effects). Realistically, no real-world signals (including SOAE) are truly stationary, but it is an important approximation crucial to many statistical assumptions. While the “system parameters” related to SOAE signals as measured in the ear canal are undoubtedly not constant, we attempted to account for such in our experimental paradigm by having subjects in the booth for some period of time (*>* 15 minutes) to adjust to the quiet environment prior to measurements, recording SOAE signals over a relatively short observation period (typically 2 minutes), and using waveforms without acoustical or electrical artifacts.

**Supplementary Figure S4:**
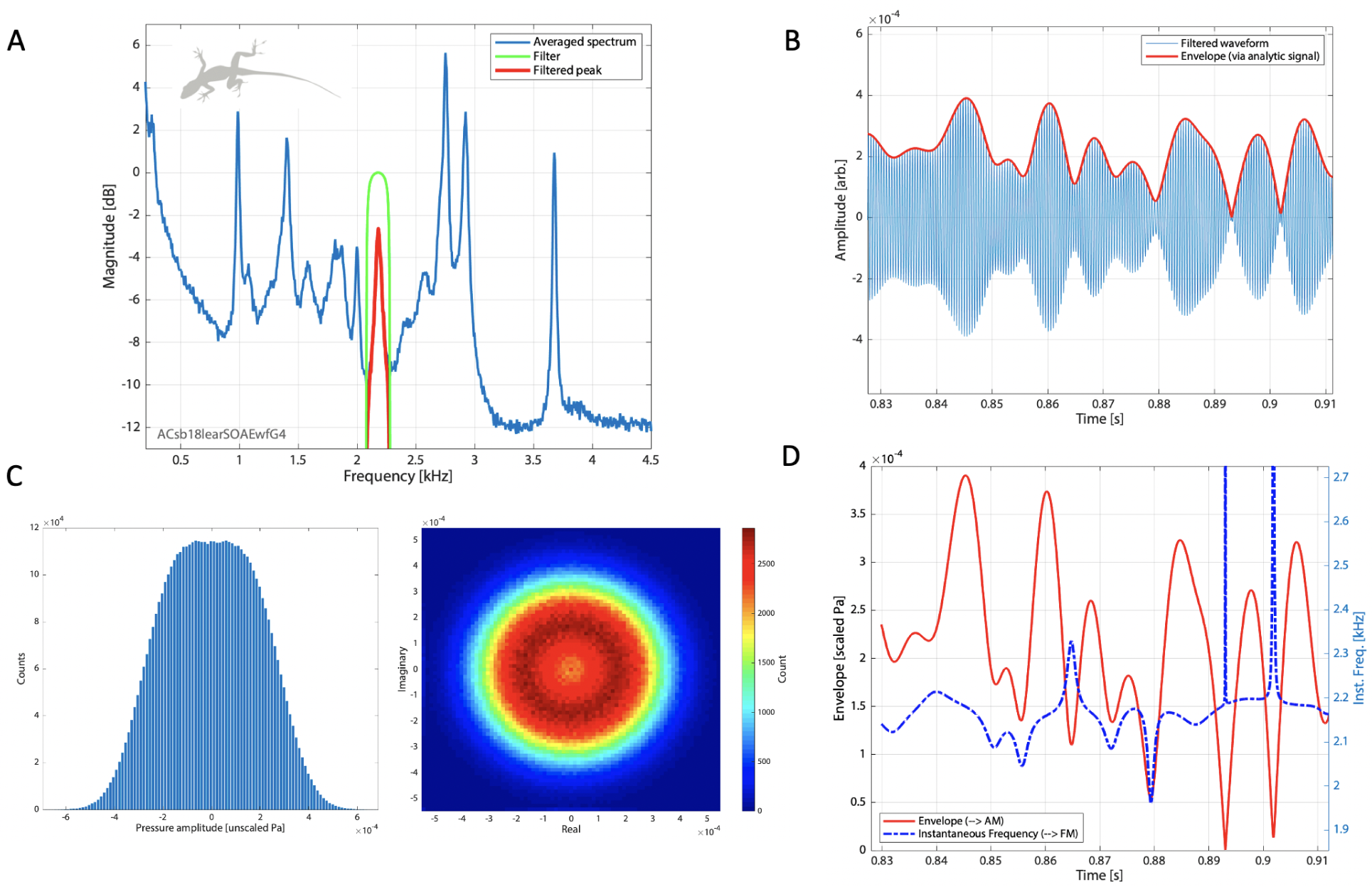
Overview of the peak-filtering process. **A** – Magnitude of SOAE spectrum from an anole lizard (blue). Green curve is the recursive exponential filter, which is multiplied to obtain the filtered signal (red). For the peak shown, the filter had a center frequency of 2.178 kHz and half-width of 100 Hz (total range span is 200 Hz). **B** – The filtered signal, including the phase (not shown), is inverse transformed to obtain the corresponding time waveform. Only a short segment is shown here for clarity. The analytic signal is then computed, from which the envelope (red curve) and instantaneous frequency (not shown) can be determined. **C** – Amplitude (left) and analytic signal (right) distributions for the peak, which visually suggest sinusoidal-like (i.e., bimodal) statistics. Note, however, that Hartigans’ dip statistic classified the amplitude distribution as unimodal. **D** – The analytic signal provides both the envelope (red solid line; identical to panel B) and instantaneous frequency (blue dashed line). These fluctuations then provide the basis for characterizing the peak’s amplitude (AM) and frequency modulations (FM), respectively.

SOAE patterns were reasonably stable across the recording time when using 46 ms windows to compute a short-time Fourier transform to observe gross temporal variations (Fig.S2). This is additionally supported by analytic signal distributions: if an emission was turning on/off, we might expect a superposition of caldera and molehill distributions, which was not generally observed (e.g., Fig.S4C). There was one obvious exception where a large peak (25 dB SNR) at 8.3 kHz in a human appeared to turn on/off (Fig.S14), similar to that previously noted for a barn owl (Fig.4A of van Dijk et al., 1996, see also Burns et al. 1984). We attempted to only analyze waveforms that did not include artifacts (acoustical or electrical), though this was not always the case for owls or humans (e.g., subject JI). When present, artifacts could be visibly discerned in the entire SOAE waveform, or as brief excursions in phase-plane plots of the ENV_p_ or IF_p_ signals (see Fig.S10). Barring an artifact, phase-plane plots typically showed qualitatively consistent properties across all filtered peak signals. Thus, we assume the measured SOAE waveforms can reasonably be assumed to be approximately stationary to first order. Nonetheless, we report several facets related to this consideration as follows. In summary, there were some deviations from stationarity that likely warrant further study, especially for humans, that may suggest richer dynamics at play in the mammalian cochlea. Those aspects will not be considered further here and the rest of the analysis detailed in the Results focuses on methodologies that assume stationarity.

**Supplementary Figure S5:**
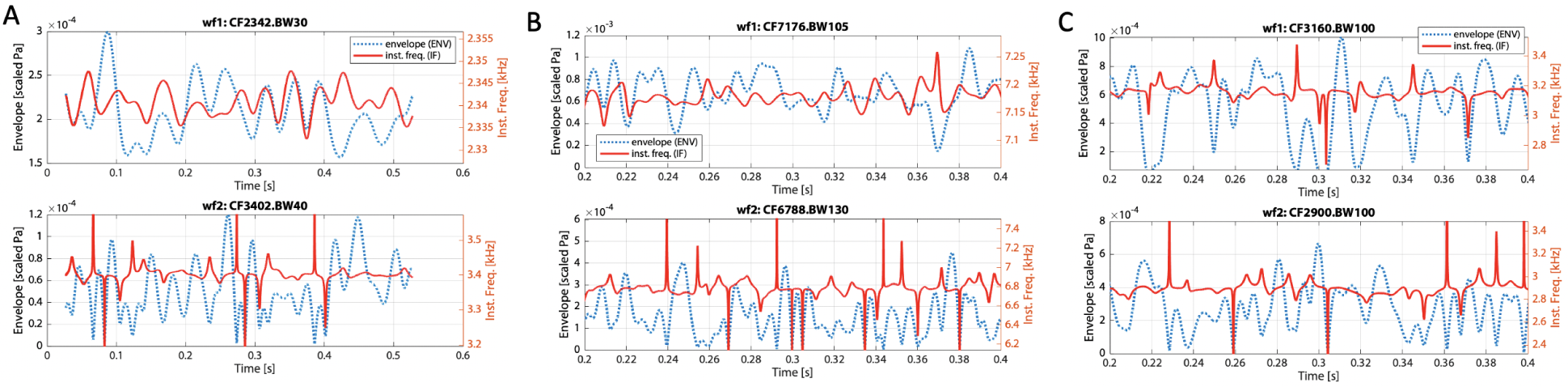
Representative amplitude modulation (AM; blue dashed lines) and frequency modulation (FM; red solid lines) curves for a pair of peaks from a representative individual from each species. For each species, the top row shows a relatively tall peak, while the bottom shows a comparatively shorter one. Peaks were picked from the same spectra shown in Fig.S3 as follows: **A** – Human (JIrearSOAEwf2) at 2.3 kHz (top, 30 Hz filter bandwidth) and 3.4 kHz (bottom, 40 Hz), **B** – Owl (TAG4learSOAEwf1) at 7.2 kHz (105 Hz) and 6.8 kHz (130 Hz), **C** – Anole (ACsb27learSOAEwfA1) at 3.2 kHz (100 Hz) and 2.9 kHz (100 Hz). Note the different timescales between plots, as well as the dual vertical axes (amplitude scale on left, instantaneous frequency on right).

### S1.5 Results: Additional Fluctuation Analyses

Additional considerations concerning the two SOAE peaks from an anole lizard shown in Fig.S6 (at 1.7 and 1.9 kHz) are described here to provide further context for the fluctuation analyses. Unless noted otherwise, the general features/shapes identified here were preserved across species independent of whether a given peak was classified as uni- or bimodal. First in Fig.S9, autocorrelations and waveform distributions for the two peaks are shown. In general, autocorrelations for AM (left column, blue curves) were relatively localized in time (i.e., a single large peak) at zero lag, though varying degrees of sinc-like behavior could be seen away from *t* = 0. The FM autocorrelations (red column, blue curves) were also temporally localized in time, but were more sharply peaked and concave-up about *t* = 0. Amplitude distributions of the AM waveforms (middle column) appeared Poisson-like, while FM distributions (right column) appeared Lorentz-like. Occasional deviations from this pattern were observed. For example, some AM autocorrelations showed a greater degree of “ringing” compared to other peaks, or a widened elevated baseline was superimposed atop. These features were considered as being indicative of some sort of non-stationary excursions and were most commonly observed for SOAE peaks from humans and, to a lesser extent, anole lizards. Second, Fig.S10 shows phase-plane portraits for the two aforementioned peaks. The entire filtered waveform (95 s) was used here. Within a given peak (i.e., between a peak’s envelope and instantaneous frequency; top row), the phase plane shows a characteristic “Christmas tree” shape centered about the CF. Across peaks, the AM (bottom left) phase plane typically appears as a filled-in rectangular region, while the FM (bottom right) appears as a plus sign. In general, while relative heights and widths could vary across peaks and species, these shapes were preserved.

Figure S11 shows a SOAE peak pair from an anole lizard with a relatively strong IPPxcAM (Fig.S11B) to illustrate the effect of averaging for the correlation analysis. As apparent in the two heat maps obtained with different averaging segment lengths (panels C and D), the correlation of the envelope fluctuations was visibly apparent as the horizontal band about 0 ms lag. While the values in panel C could be as high as 0.6 at any one time point in the band, their average was approximately 0.16, corresponding to the peak in the curve at 0 ms lag in panel B. Typically, however, most correlations were weaker (correlation coefficient *<* 0.1) and the corresponding heat maps did not show visually distinguishable bands indicative of a correlation. That is, averaging was necessary to extract the correlation.

**Supplementary Figure S6:**
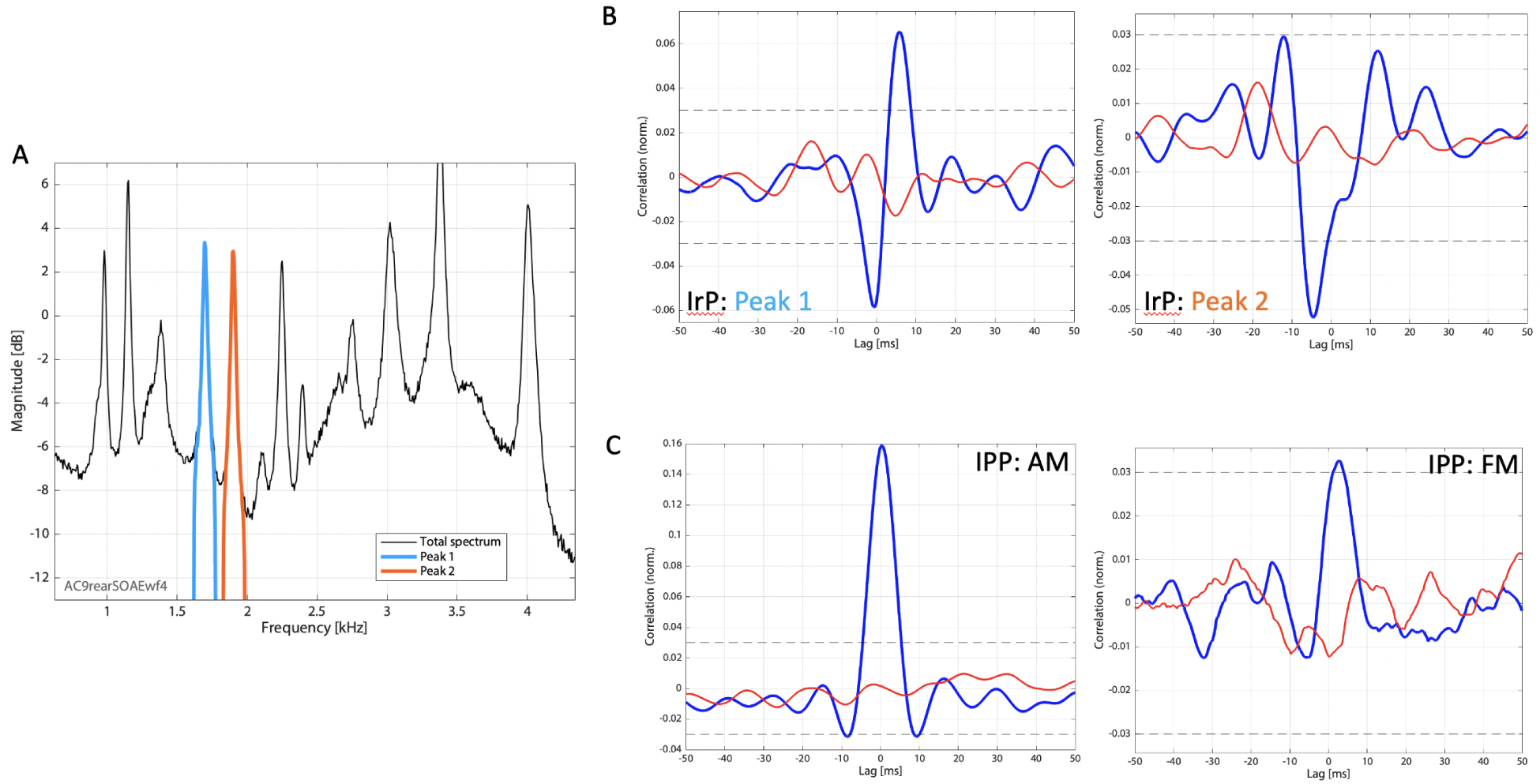
Overview of the correlation method and illustrative results. **A** – Representative SOAE spectra from anole lizard (AC9; same as bottom panel in Fig.1) with two peaks isolated at 1.7 and 1.9 kHz (blue and orange, respectively) for comparison (see Fig.S4). The convention used is that peak 1 is always the lower frequency peak of the pair. **B** – For a given peak, the “intrapeak” (IrP) correlation *T_xc_* of AM relative to FM (i.e., IrPxc) is shown in blue. The control curve *T_b_* (red) was obtained by bootstrapping (see Methods). The horizontal black dashed lines indicate the thresholds used to determine if a correlation was present. The correlation for peak 1 was deemed present but indeterminate (i.e., no single dominant peak in the IrPxc), such that neither correlation sign nor delay were determined. The correlation for peak 2 was present, normal (i.e., single dominant peak), negative (i.e., downward peak), and with a delay of −4.6 ms (negative here means that FM is delayed relative to AM). **C** – Interpeak (IPP) correlations for both AM (left; IPPxcAM) and FM (right; IPPxcFM). Positive, normal correlations were deemed present for both peaks, although this was weaker for the IPPxcFM. Further, positive delays (i.e., the lower frequency peak was delayed relative to the higher one) were observed.

In Fig.S12, additional IrP and IPP correlations are shown for an anole lizard. The top row shows IrPxc curves (i.e., AM and FM cross-correlations), whereas the bottom row shows the comparison between two peaks for their AM relative to the FM of the other. In all cases shown here, a correlation would be deemed present, however be designated “indeterminate” (as they are not isolated singular peaks). In a general sense, these correlations were considered atypical correlations due to the lack of a single peak, as was more commonly observed when an IPP correlation could be discerned. Although the majority of IrPxc were not singly-peaked, these were likewise considered atypical for consistency. For additional context, Fig.S13 demonstrates several shapes that arose when a noisy waveform was cross-correlated with its various time derivatives. Numerous features from the SOAE data shown in Fig.S12 showed similar structure.

A different way to visualize the filtered waveform distributions is shown in Fig.S14. To show the time evolution of the signal, SOAE peaks from a human subject (JI) were filtered and shorter segments (190 ms here) were binned as histograms and plotted sequentially as a heat map. The 2.3 kHz peak in panel B clearly shows a strongly bimodal nature and was classified as such, while the 5.8 kHz peak in C was classified as unimodal (albeit displaying bimodal properties visually). The most striking feature here is in panels D and E, where both peaks appear to dramatically change their behavior: the 8.3 kHz peak shifts between bimodal and unimodal distributions, whereas the 8.7 kHz peak appears to turn off. Changing filter BW did not affect this behavior. This sort of dynamic was only noticed very rarely, though may be consistent with results previously reported for a barn owl (van Dijk et al., 1996). Further, there were two small “sideband” peaks, a small one approximately 377 Hz below and a slightly larger one 366 Hz above, the latter being strongly AM-correlated to the central peak whereas the former was not.

**Supplementary Figure S7:**
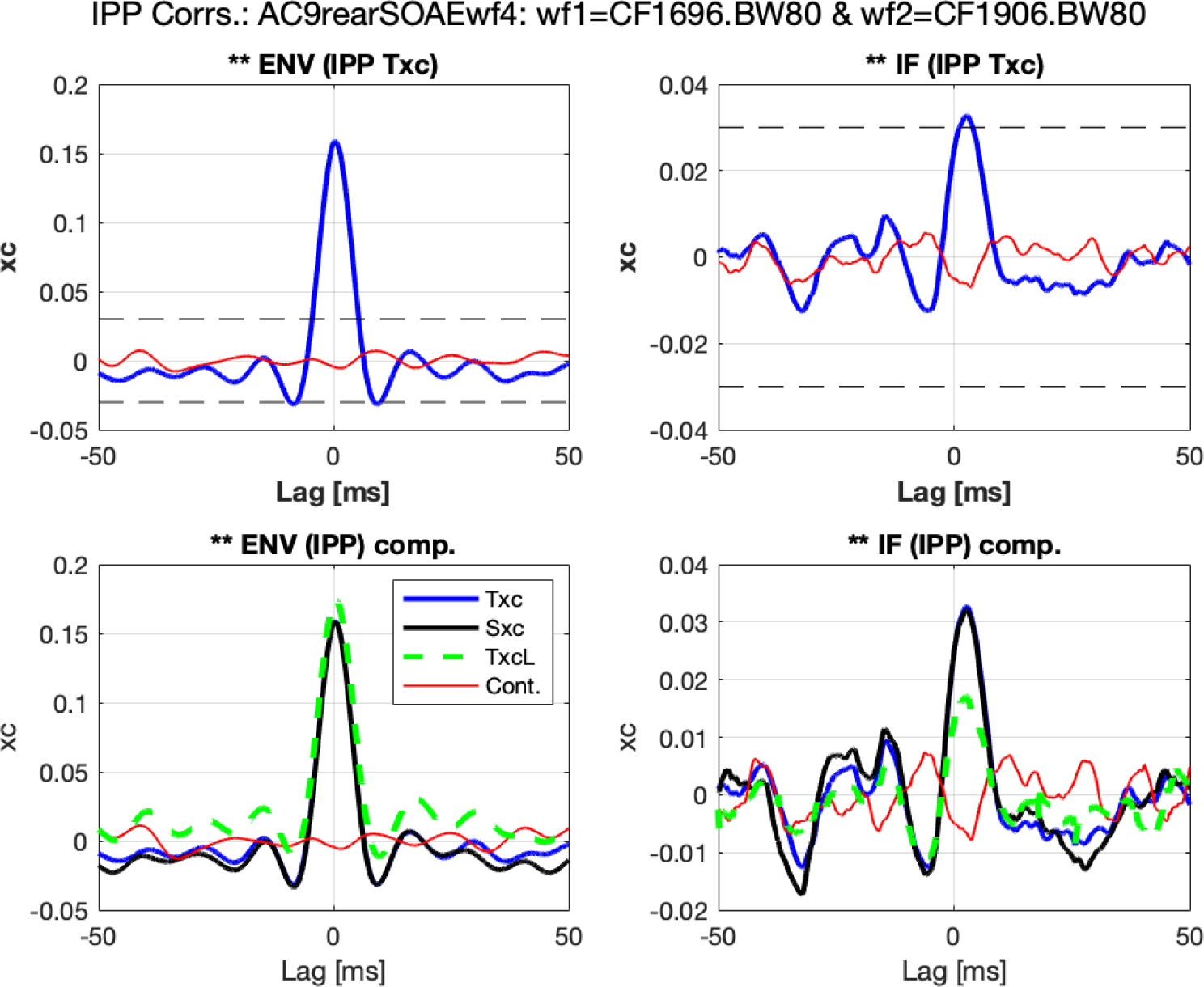
Comparison of IPPxcAM and IPPxcFM correlation curves for the same two peaks shown in Fig.S6 from an anole lizard (AC9, wf1= 1.696 kHz peak and wf2= 1.906 kHz). The top row shows the correlation curves obtained from the short-duration averaged time-domain calculation (*T_xc_*) that was the basis used for the values reported in the main body of the paper. The bottom row shows same curves as the top row (blue solid line) compared against correlation curves computed via the time-domain for the entire waveform (green dashed line) and the short-duration averaged spectral-domain calculations (black). By and large, similar results are obtained by all three approaches. All correlation values were normalized to be between [*−*1, 1].

### S1.6 Results: General Comparison of AM Correlations Across Species

Figure S16 shows a visual comparison across species of peak height and width (via the Lorentzian fits as described in the Methods) versus frequency relative to whether a given peak pair showed AM correlation (IPPxcAM) and/or were neighbors. In general, IPPxcAM occurred between SOAE peaks that were taller and narrower in barn owls. SOAE peak heights and widths were not related to the presence of IPPxcAM for humans and anole lizards. Further, Fig.S16 shows that IPPxcAM was more likely to be present for peaks that were nearest-neighbors (NNs) in barn owls and anole lizards, but not humans.

Figure S17 shows a comparison of interpeak amplitude modulation cross-correlation (IPPxcAM) curves for a peak representative pair from each of the three species.

**Supplementary Figure S8:**
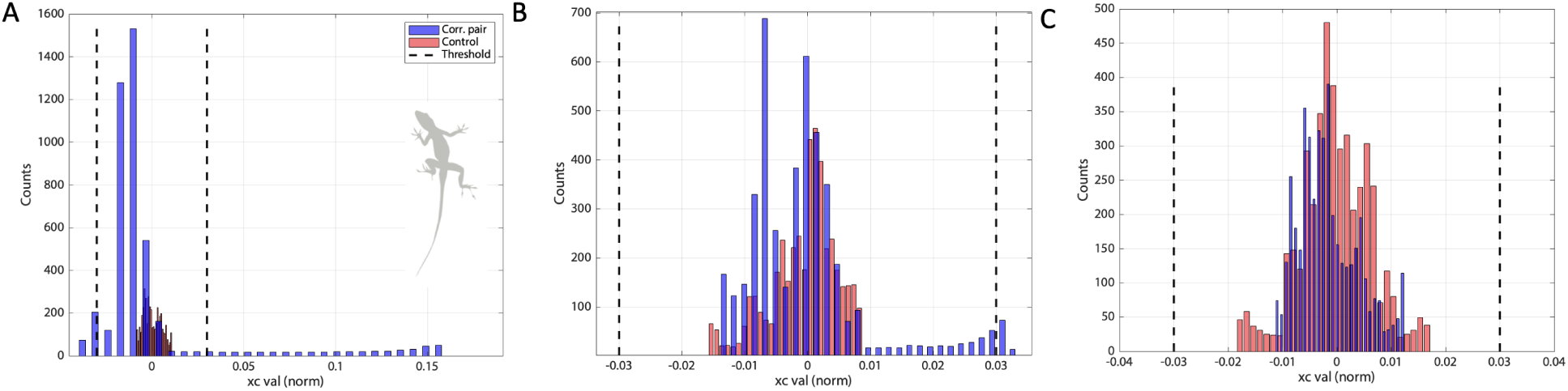
Histograms for (normalized) interpeak cross correlation values as shown in top left panel of Fig.S7 for an anole lizard (AC9). Values for the pair in question are in blue and those for the (randomized time-shifted) control pair are in red. Vertical dashed lines show the threshold criterion bounds for classification as to whether a correlation was present. **A** – The time-domain IPPxcAM for wf1 = 1.696 kHz peak and wf2 = 1.906 kHz. **B** – Same pair as **A**, but for the IPPxcFM. **C** – The IPPxcAM for two peaks not exhibiting any correlation (wf1 = 1.696 kHz peak and wf2 = 2.865 kHz).

**Supplementary Figure S9:**
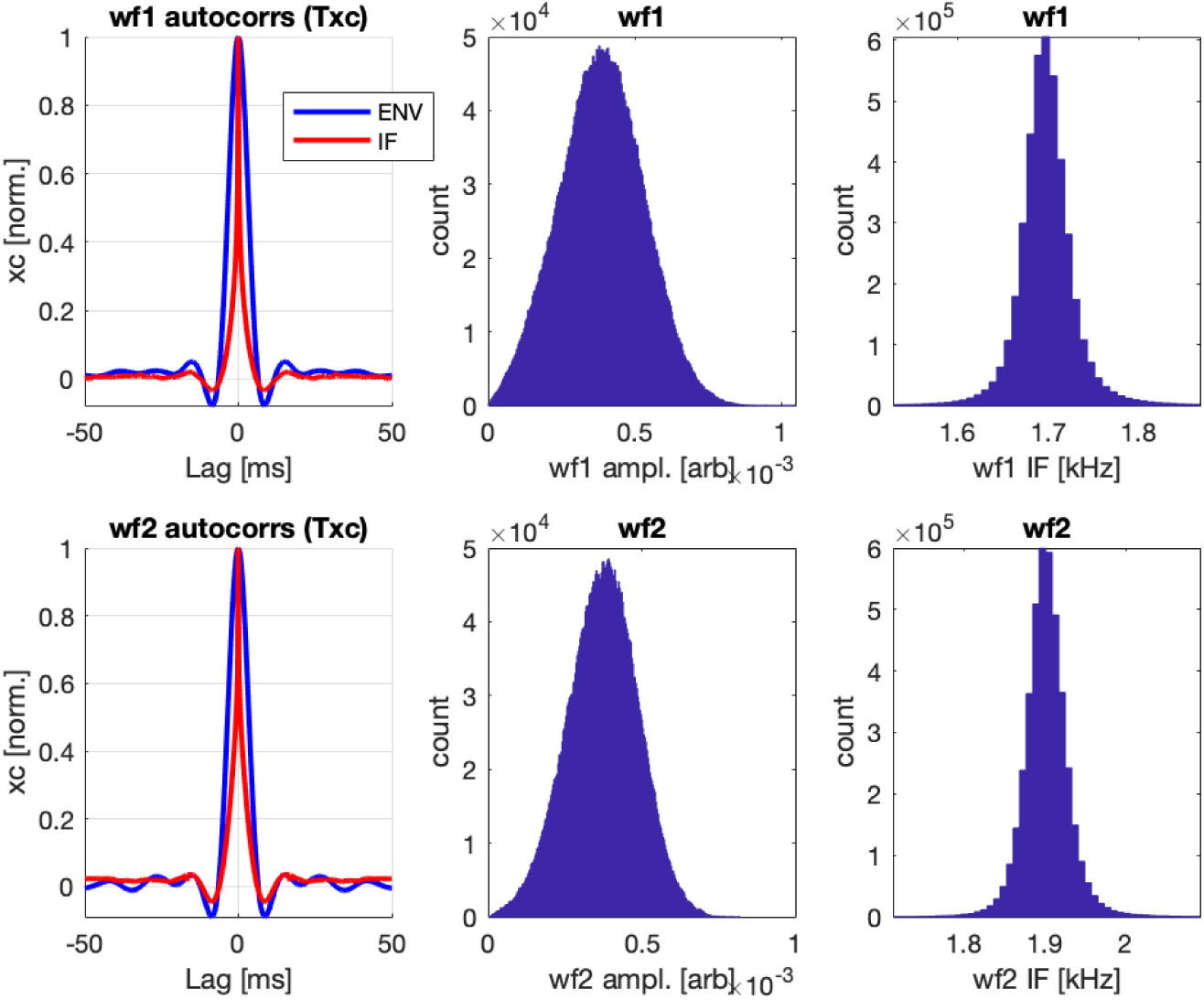
Intrapeak (IrP) relationships for the two SOAE peaks shown in Fig.S6 from an anole lizard (AC9). Top row: wf1= 1.696 kHz peak, bottom row: wf2= 1.906 kHz. Left column shows the averaged autocorrelations for the envelope (AM, blue) and instantaneous frequency (FM, red), computed similarly to the cross correlations described in the main document. AM (middle column) and FM (right) distributions were obtained from the entire (95 s) filtered waveforms.

### S1.7 Results: Additional Considerations

Several additional facets relevant to the results but not systematically explored in this study are briefly described below.

#### S1.7.1#Factors Affecting Correlative Behavior

One consideration is how stable the relations shown in these correlation maps are across time, beyond the 1-2 minute SOAE waveform acquisition time. For example, how interpeak relations may change in the few hours over which an experiment takes places (also in light of perturbation from external acoustic stimuli), or days/years over different experiments. Homeostatic conditions (e.g., body temperature) need to be accounted for to properly assess such. However, one anecdotal measurement is presented here, as it also addresses the relatively rare occurrence of interpeak FM correlations (e.g. there is only one instance in Fig.4; bottom right map just below 2 kHz). It deals with an anole lizard whose SOAE is shown in the top right panels of Figs.1 and 4. For that experiment, the SOAE waveform chosen for analysis occurred early in the experiment after body temperature stabilized. This animal had been slated for euthanasia for veterinary reasons, though exhibited relatively strong SOAE activity, which remained strong throughout the experiment. Several hours after the primary waveform was recorded, the animal was given a large overdose of anesthetic with the expectation that SOAE activity would disappear, as has been observed in the past. However, the converse was observed: the SOAE activity increased and showed relatively enhanced correlative behavior, as shown in Fig.S20. The downward frequency shift apparent is likely tied to body/head temperature changes and/or change in physiological state. While the animal showed no visible respiration at the time of measurement, no other diagnostic measures of physiological state were made prior to decapitation, so presumably the animal was only deeply anesthetized. Note that the correlation map in Fig.S20 shows significantly more instances of interpeak FM correlations, though sometimes required a lower statistical threshold for inclusion. Upon decapitation, all SOAE activity disappeared.

**Supplementary Figure S10:**
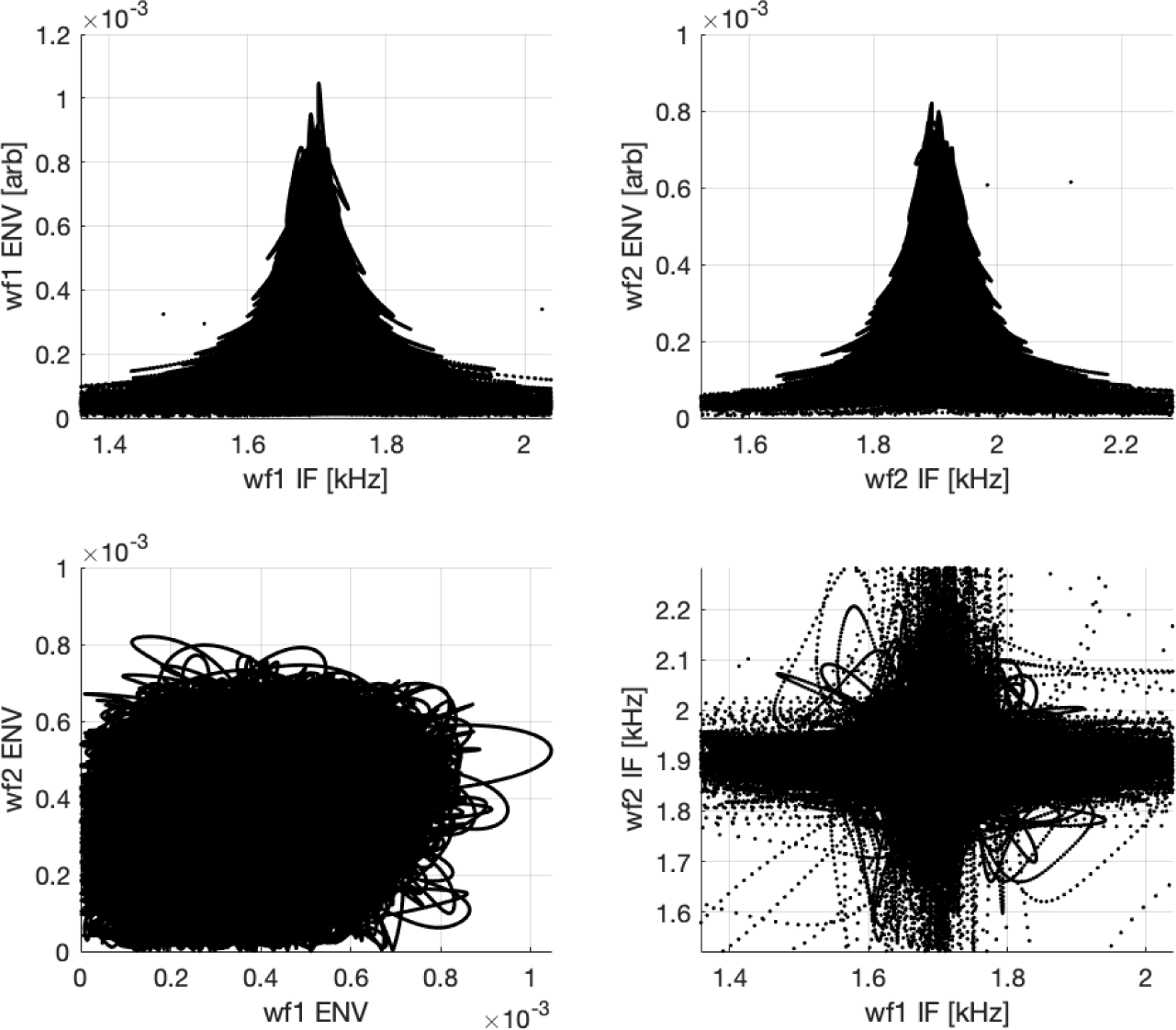
Phase-plane relationships for the two peaks shown in Fig.S6 from an anole lizard (AC9). Top row shows the intrapeak (IrP) phase-plane relationship between a given peak’s envelope (AM) and instantaneous frequency (FM) fluctuations (wf1= 1.696 kHz peak on left, wf2= 1.906 kHz on right). Bottom row shows the interpeak (IPP) phase-plane relationships for the envelope (left) and instantaneous frequency (right).

Another consideration is how SOAE correlations may be affected by perturbations due to external stimuli. It has been well established that SOAE readily interact with tones, and that such relates to evoked emissions (e.g., Bergevin et al., 2012, 2015). To illustrate this, Fig.S21 shows several spectrograms for swept tones and/or tone bursts for an anole, and similar results are observed for human and owl (Bergevin and Salerno, 2015). In short, SOAE activity is affected by stimuli in a temporally and spectrally confined fashion (e.g., vertical bands for the tone bursts in panels C & D). General SOAE spectral patterns are, however, stable to these perturbations, and thus one might expect correlative patterns to be as well. However, data from at least one ear indicate it is unclear to what extent this holds. For one anole (ACsb24), SOAE waveforms were collected about an hour apart (files A1 and B1). The spectral structure remained fairly stable, albeit with a slight upward frequency shift. During the hour between, various SFOAE runs using moderate-level swept tones were presented. The general correlative features largely remain in place, although some minor changes are apparent (e.g., shift in delays, disappearance/emergence of some IPPxcFM). This observation suggests that while general spectral structure might be fairly stable, interpeak correlative relations might vary more fluidly with time and/or in response to external stimuli.

**Supplementary Figure S11:**
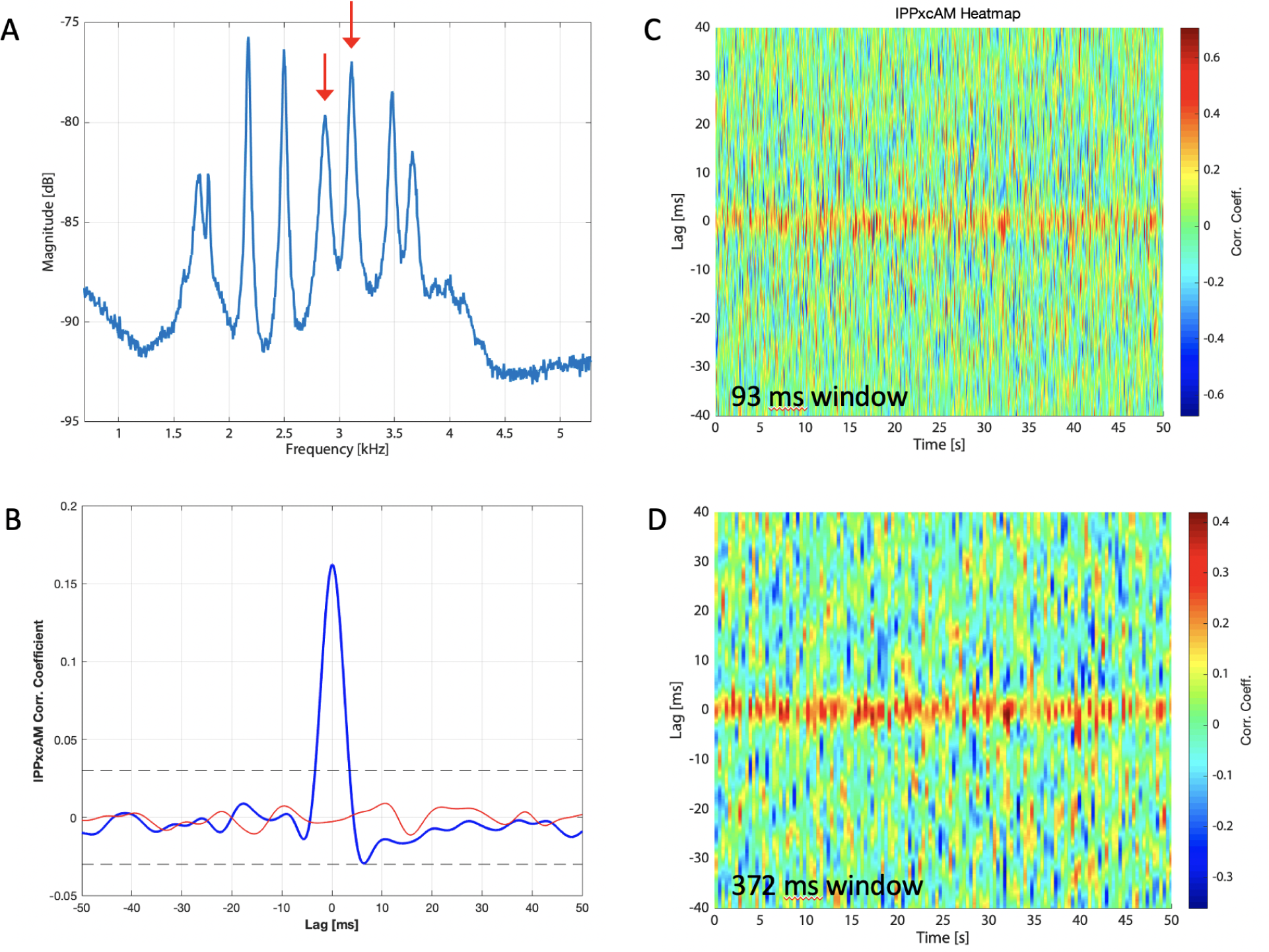
Considering the correlation as a heat map. SOAE data from an anole (ACsb24). **A** – Averaged spectrum, with two peaks (red arrows at 2.9 and 3.1 kHz) chosen for IPPxcAM analysis. **B** – Associated averaged IPPxcAM curve, showing a suprathreshold correlation response. **C** – Heat map showing the individual correlation curves that were averaged across time to obtain the curve in panel B. Here, the segment window was 4096 points long (93 ms long). **D** – Same as panel C, but a longer window of 16384 points per segment (372ms) was used.

#### S1.7.2#Non-peak Correlations: Valleys Between

As described briefly in the main text, anecdotal correlation analyses for the “valleys” between peaks is shown for one anole in Fig.S18. While no FM correlation was observed, a weak positive AM correlation was observed for this pair. This indicates correlations for valleys can also manifest, presumably related to underlying SOAE “baseline” activity (Manley et al., 1996).

As noted prior, a feature of SOAE activity that appears unique to lizards was the presence of “baseline” activity (Köppl and Manley, 1993; Manley et al., 1996; Manley and Gallo, 1997), as visually apparent in Fig.1. This baseline activity is also suppressible in that it readily interacts with external sounds (e.g., Fig.S21). Given high vector-strength values in SOAE activity across ears at non-peak frequencies (Roongthumskul et al., 2019), one might also consider the possibility of correlations for the valleys between peaks. Preliminary observations for one anole (considering the same SOAE spectrum as shown in the bottom right panel of Fig.1) are shown in Fig.S18. This figure indicates that such correlations can in fact be observed. There, both filtered signals exhibited unimodal amplitude distributions, and showed no FM correlation. However, a weak positive correlation was observed that had a negative delay (i.e., the higher frequency region lagged behind the lower one). Thus, correlative behavior need not exclusively be confined to peaks. While not systematically explored, it appears this sort of correlative behavior occurs due to the presence of the broad baseline SOAE in anoles and not due to noise in the microphone signal unrelated to SOAE.

**Supplementary Figure S12:**
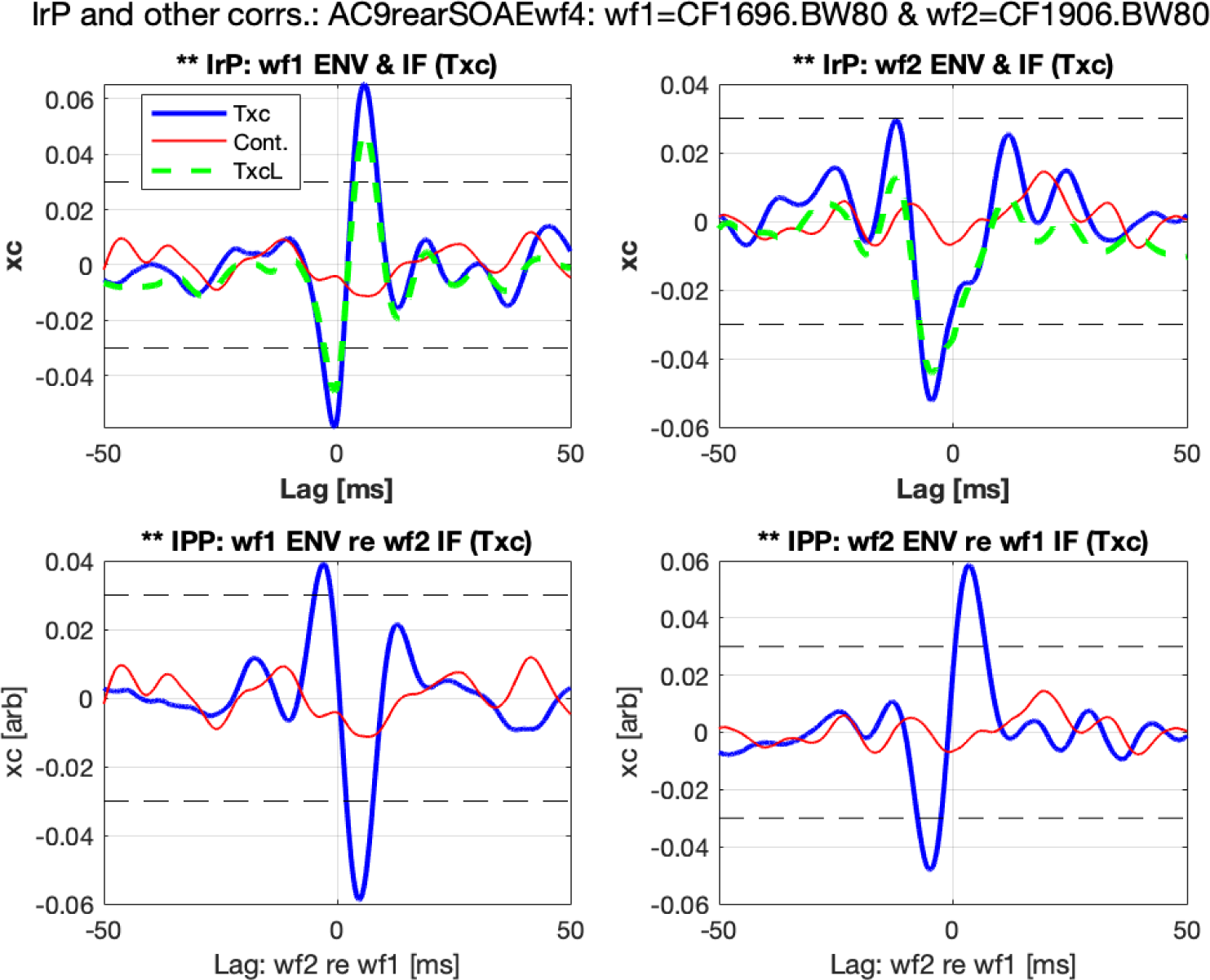
Same two peaks shown in Fig.S6 from an anole lizard (AC9, wf1= 1.696 kHz peak and wf2= 1.906 kHz). Top row is similar to the bottom row of Fig.S7, although for intrapeak (IrP) correlations (e.g., top left shows wf1 AM and FM correlation). Bottom row shows IPP correlations associated wf1 AM and wf2 FM (left) as well as wf1 FM and wf2 AM (right).

#### S1.7.3#Consideration of Other Lizard Groups

Papillar morphology varies significantly across lizard families (e.g., number of hair cells/ear, structure/presence of tectorial membrane; see Wever, 1978; Miller, 1981; Manley, 2000), motivating application of the correlation methodology employed here to study SOAE interpeak correlations for other lizard groups. While the SOAE data for lizard presented in this paper thus far focus solely on *Anolis*, an iguanid with a small basilar papilla containing hair cells whose bundles are predominantly free standing, the correlation analyses described here have been preliminarily applied to other lizard species. One example for a tegu, a teeid lizard with a larger basilar papilla whose hair cells are covered by a continuous tectorial membrane, is shown in Fig.S19. SOAE activity from these ears tend to exhibit a broad baseline with several narrow large peaks (Manley et al., 2018). The two large peaks identified in Fig.S19A showed relatively strong and complex correlative behavior (panels B-G), including robust IPPxcFM.

**Supplementary Figure S13:**
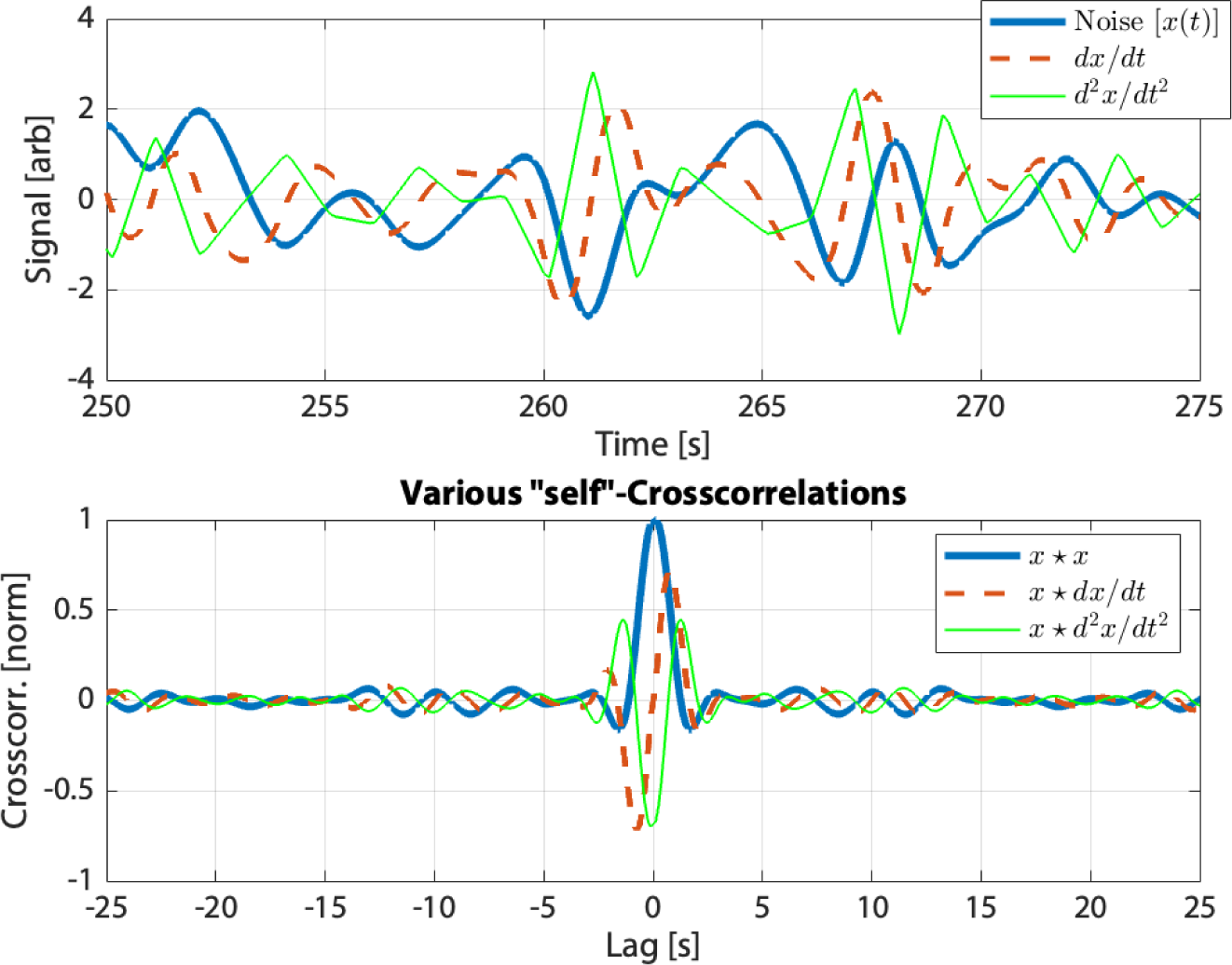
Top panel shows a low-pass filtered noisy waveform *x*(*t*) (thick blue line) and two associated time derivatives. Bottom panel shows various (normalized) crosscorrelations: *x * x* (i.e., the autocorrelation, thick blue line, *x * dx/dt* (red dashed), and *x * d*^2^*x/dt*^2^ (thin green).

Preliminary fluctuation correlation analysis was performed on SOAE waveform data collected in other lizard species. One illustrative example is shown in Fig.S19, where the SOAE activity was collected from a sub-adult black and white tegu lizard (*Salvator merianae*, formerly classified as *Tupinambis teguixin*). Some previous reports have considered Tupinambis SOAE activity (e.g. (Manley et al., 2018)), as well as SOAE peak correlations (van Dijk et al., 1998). According to (Wever, 1978), this species has a papilla length of about 1.4 mm, approximately 1400 hair cells total, all of which are covered by a continuous tectorial membrane. Here, a 60 second waveform was collected in a similar fashion to that described in the main text. The two largest peaks (as identified in Fig.S19A) both exhibited bimodal distributions. As apparent in panels B–G, strong correlative behavior was observed, although it was not well localized in time (i.e. the majority of the correlations would be classified as non-singular).

#### S1.7.4#Changes Across A Given Experimental Session & Effects of Anesthetic Overdose

Results from one Anolis experiment where data were obtained over a long experimental session (several hours) are shown in Fig.S20. In this case, the data were obtained after the deeply anesthetized lizard had been given a Nembutal overdose expected to euthanize it and body temperature had partially restabilized. However, unexpectedly, robust SOAE activity remained until decapitation occurred. These data provide some preliminary insight into how fluctuation activity may vary across a given experiment, as well as with physiological changes in the anesthetized state.

### S1.8 Response to External Swept Stimuli

Examples of spectrograms of Anolis SOAEs in responses to frequency-swept external tonal stimuli are shown in Fig.S21. Here, SOAE peaks appear as horizontal lines, whereas the stimuli can be seen as the diagonal. Interactions between SOAE and the stimuli can readily be observed (e.g., the vertical bands for the tone-burst stimuli in panels C and D). Additional information can be found in (Bergevin and Salerno, 2015).

**Supplementary Figure S14:**
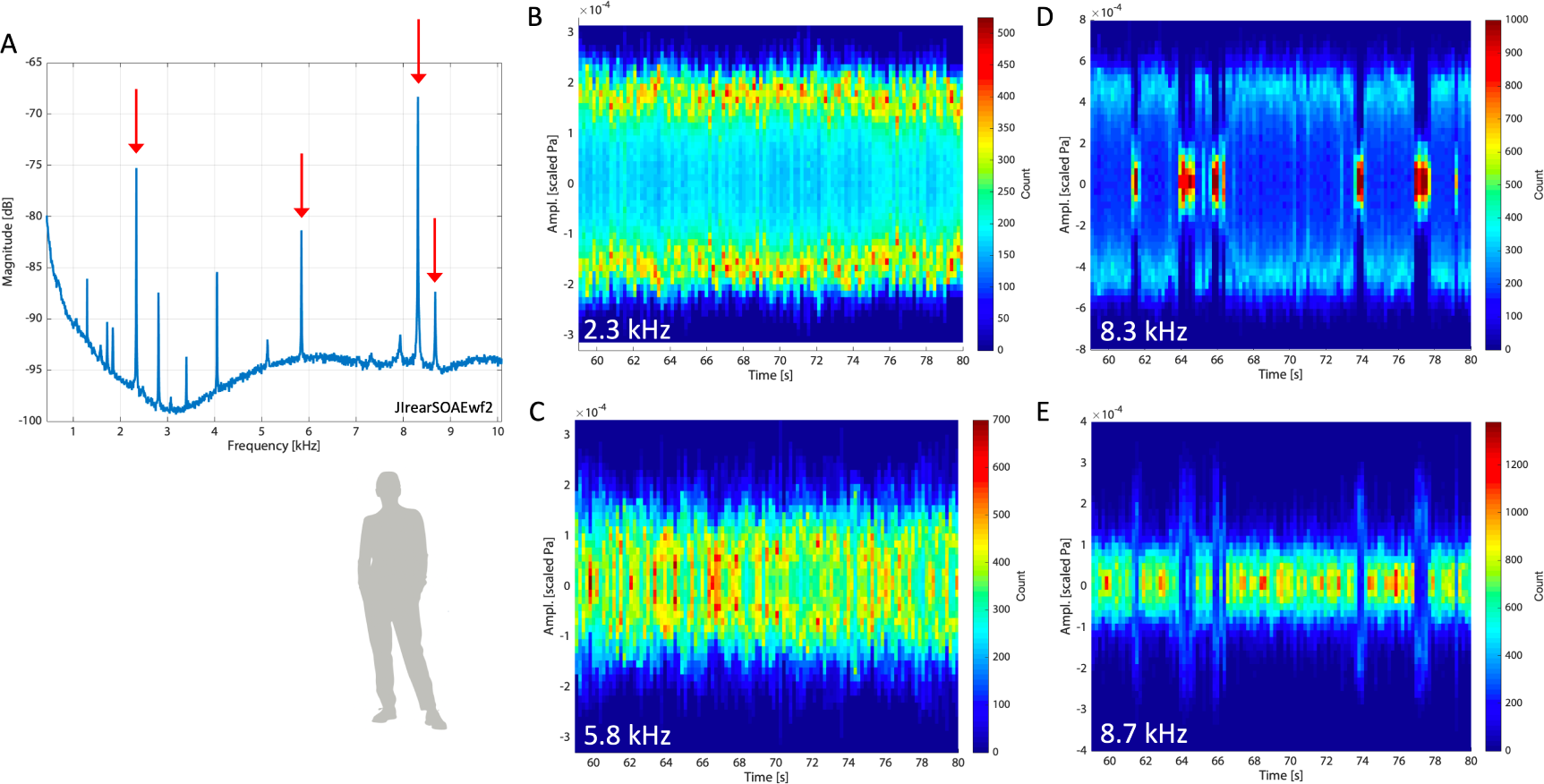
Filtered peak waveform amplitude distributions as heatmaps. **A** – SOAE spectrum from a human subject (JI). Four different peaks are marked with arrows (2.3, 5.8, 8.3, and 8.7 kHz). These are subsequently indicated in panels B–E (see SI text). The peaks shown in panels B and D were classified as bimodal via Hartigans’ dip statistic (see main Methods), whereas the peaks in C and E were unimodal. Further, the peaks shown in panels D and E (i.e., 8.3 and 8.7 kHz) were determined to exhibit IPPxcAM (but not IPPxcFM), and that the correlation was negative with a positive delay of 1.8 ms (i.e., the higher frequency peak led). A weak, indeterminate IPPxcAM was observed between the peaks at 2.3 and 8.3 kHz. No other IPP correlations were observed between these four peaks. Note that the coarser timescales shown, spanning about 20 s, are matched across all four responses so features in one can be directly compared to the others (e.g., event at 64.2 s mark in both D and E).

**Supplementary Figure S15:**
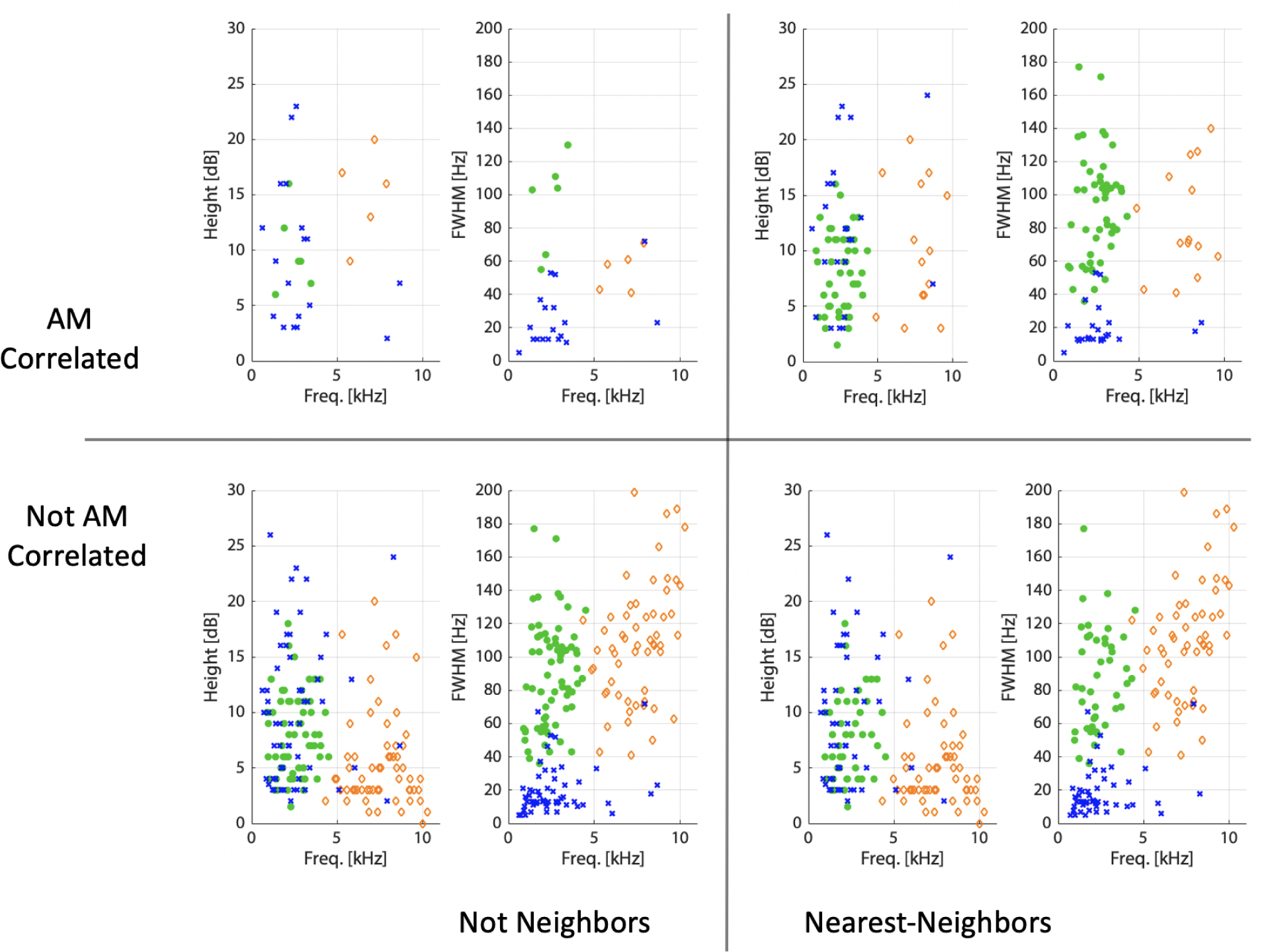
Comparison of peak height and width (as per the Lorentzian fits) across species with respect to AM correlation and relative frequency spacing. Here, peaks are categorized into one of four groups: *top left*) peak pairs exhibiting AM correlation but are not neighbors (e.g. there is at least one other analyzed peak inbetween them), *top right*) pairs exhibiting AM and are nearest-neighbors (NNs), *bottom left*) pairs not exhibiting AM correlation and not NNs, and *bottom right*) pairs not exhibiting AM correlation and are NNs. Values from both peaks are plotted, which leads to some overlapping of points (e.g. the density of points in the bottom row is actually greater than the top row than visually suggested) and there is also overlap of some values across different plots. Same legend as other figures applies (blue x – human, orange diamond – barn owl, green * – anole).

**Supplementary Figure S16:**
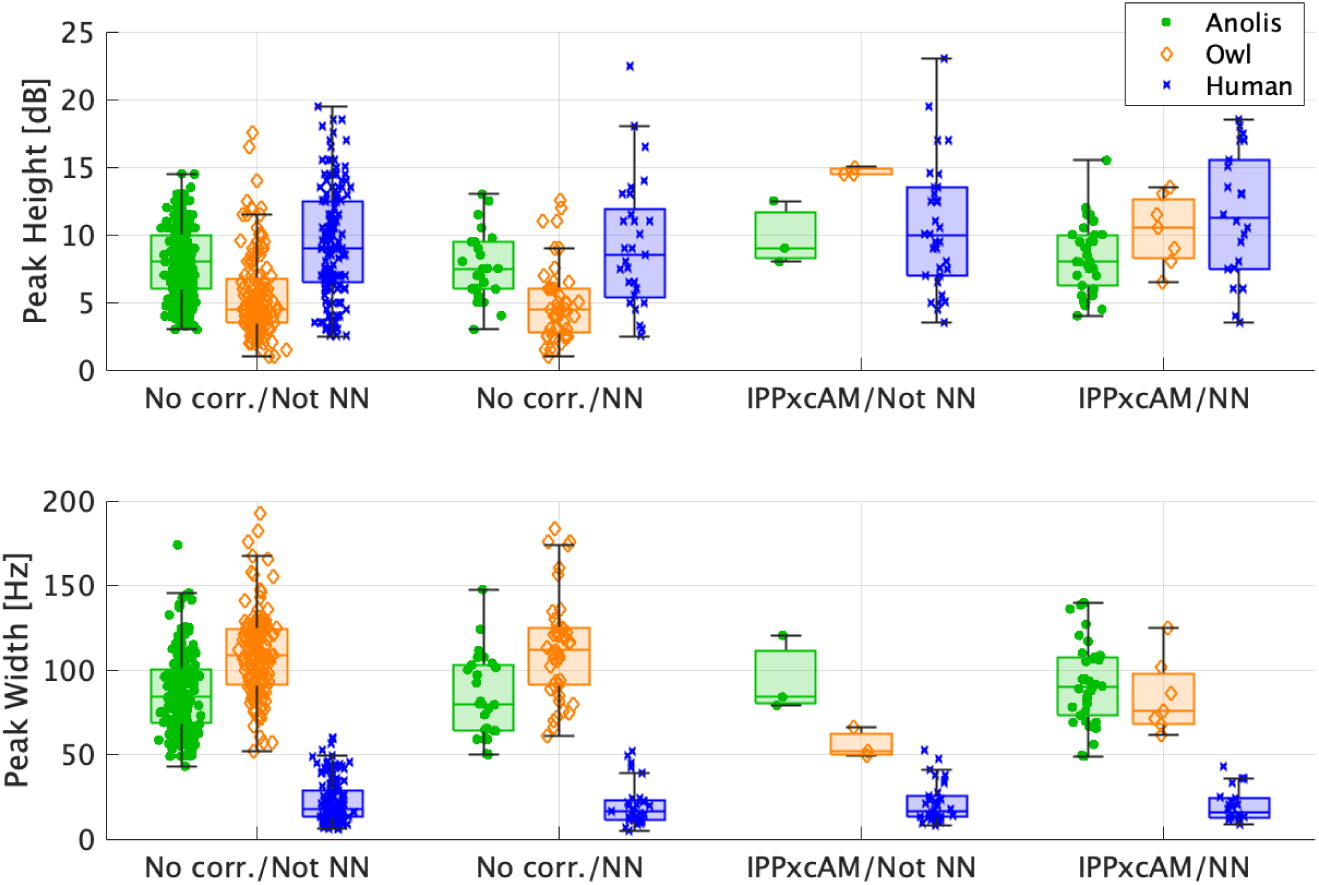
Comparison of peak pair height and width (as per the Lorentzian fits) across species with respect to the presence of AM correlations (IPPxcAM) and relative frequency spacing (peaks that were nearest neighbor pairs [NN] or not). Heights (top panel) and widths (bottom panel) are presented as the mean values for a given pair. Here, peaks are categorized into one of four groups: “No Corr./Not NN” = peak pairs without IPPxcAM that were not NN (there was at least one other SOAE peak between them), “No Corr./NN” = peak pairs without IPPxcAM that were NN (no SOAE peaks between them), “IPPxcAM/Not NN” = peak pairs exhibiting IPPxcAM but were not NN, and “IPPxcAM/NN” = peak pairs exhibiting IPPxcAM that were NN. Each box represents the interquartile range with a central mark at the median. Values beyond the boundaries of the whiskers were considered outliers. Horizontal jitter was added to improve visualization.

**Supplementary Figure S17:**
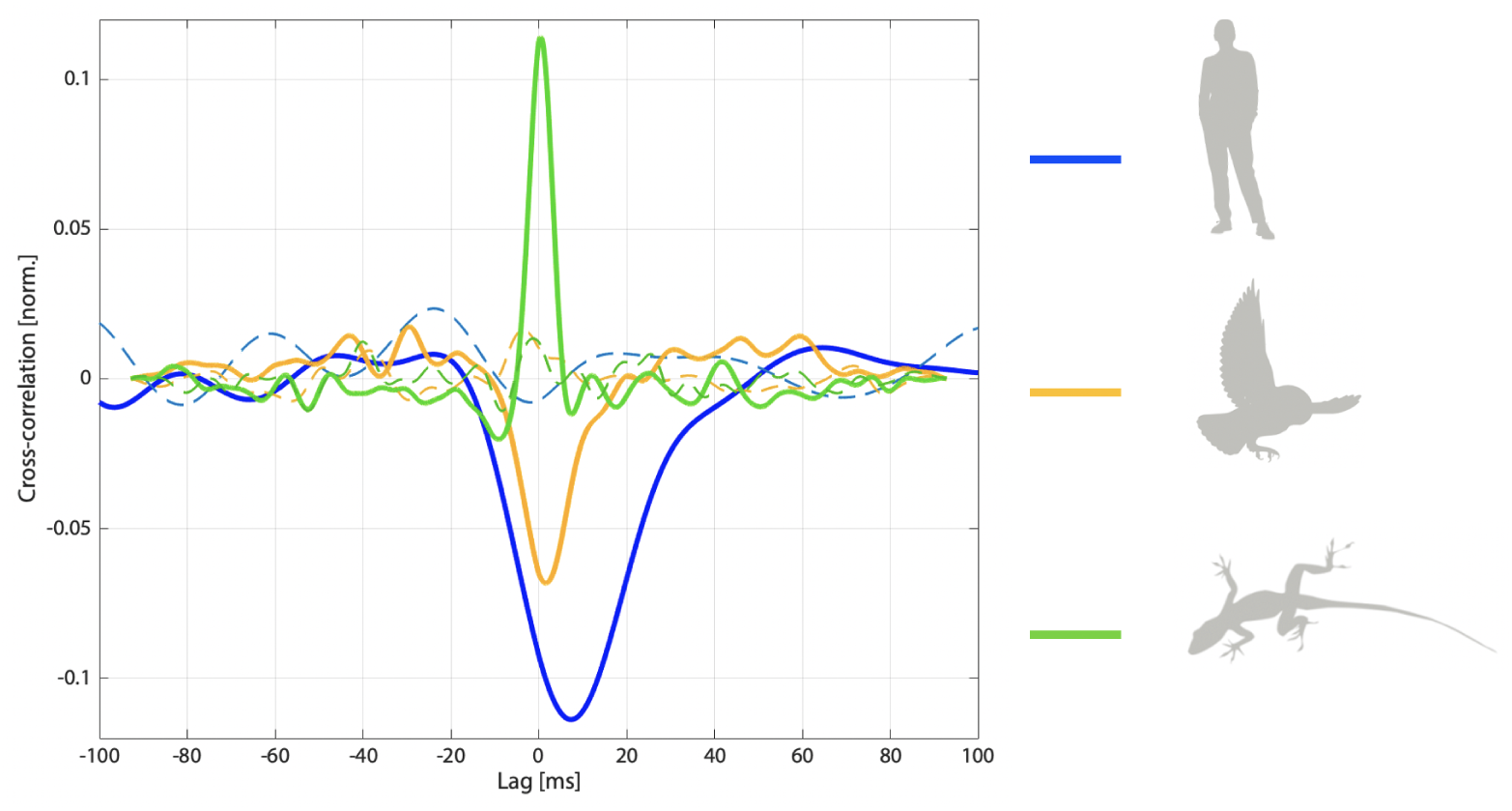
Comparison of exemplar interpeak amplitude modulation cross-correlation curves (IPPxcAM) from each species. The subjects and peaks used are as follows (with frequency in kHz and amplitude distribution classification): human (blue) = JI – 2.3 (bimodal) & 2.8 (unimodal), barn owl (orange) = TAG4 – 5.8 (unimodal) & 7.2 (bimodal), anole lizard (green) = ACsb24 – 2.5 (bimodal) & 2.9 (unimodal).

**Supplementary Figure S18:**
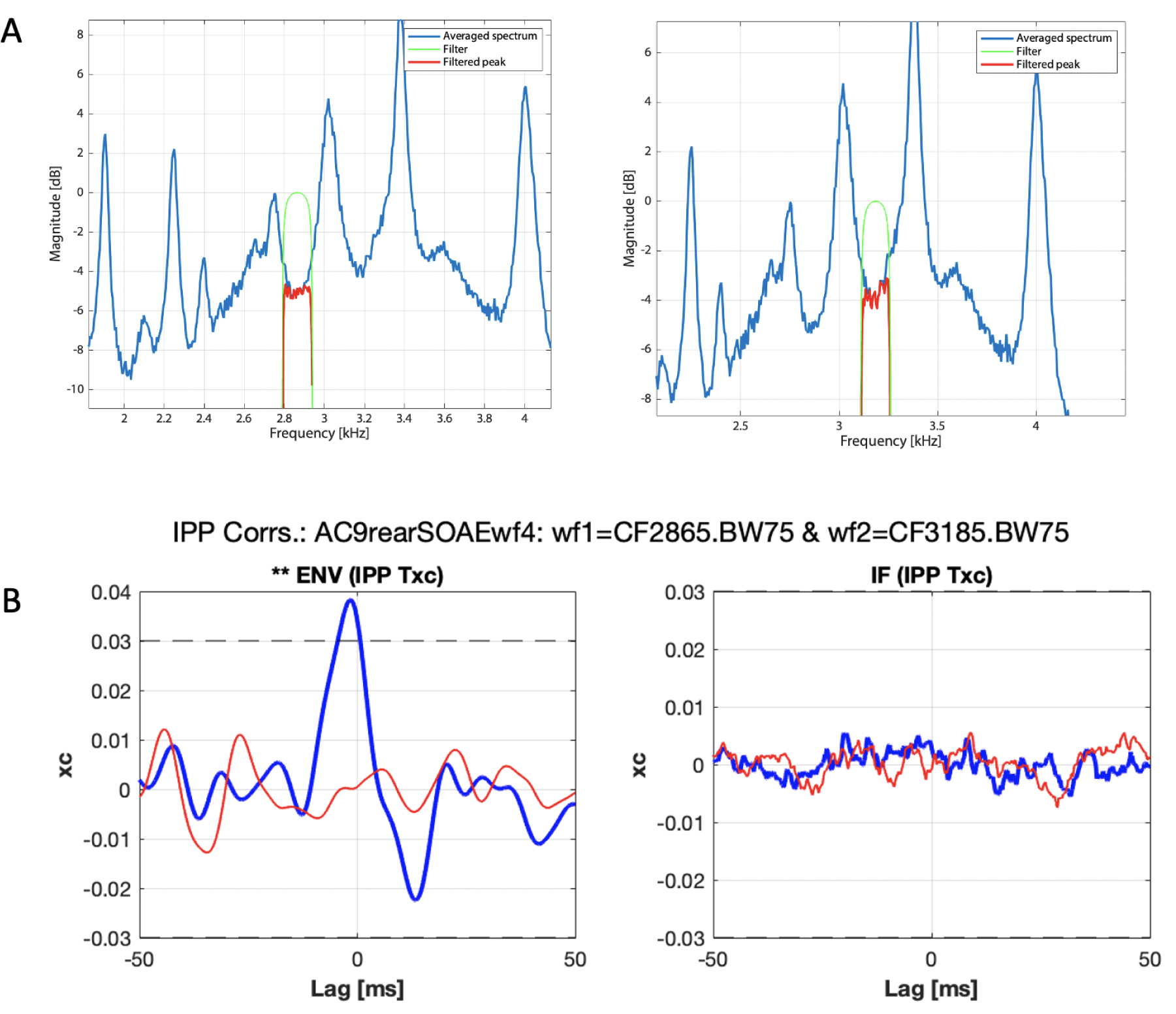
Correlation analysis applied to filtered frequency regions between peaks for one Anolis lizard. Note that this SOAE spectrum is the same one as shown in the bottom right plot of Fig.1 (AC9). **A**-Spectra for both filtered signals, with center frequencies of 2865 and 3185 Hz respectively, both with filter bandwidths of 75 Hz. **B**-Corresponding correlation curves. The AM curve on the left shows a weak, but present, correlation.

**Supplementary Figure S19:**
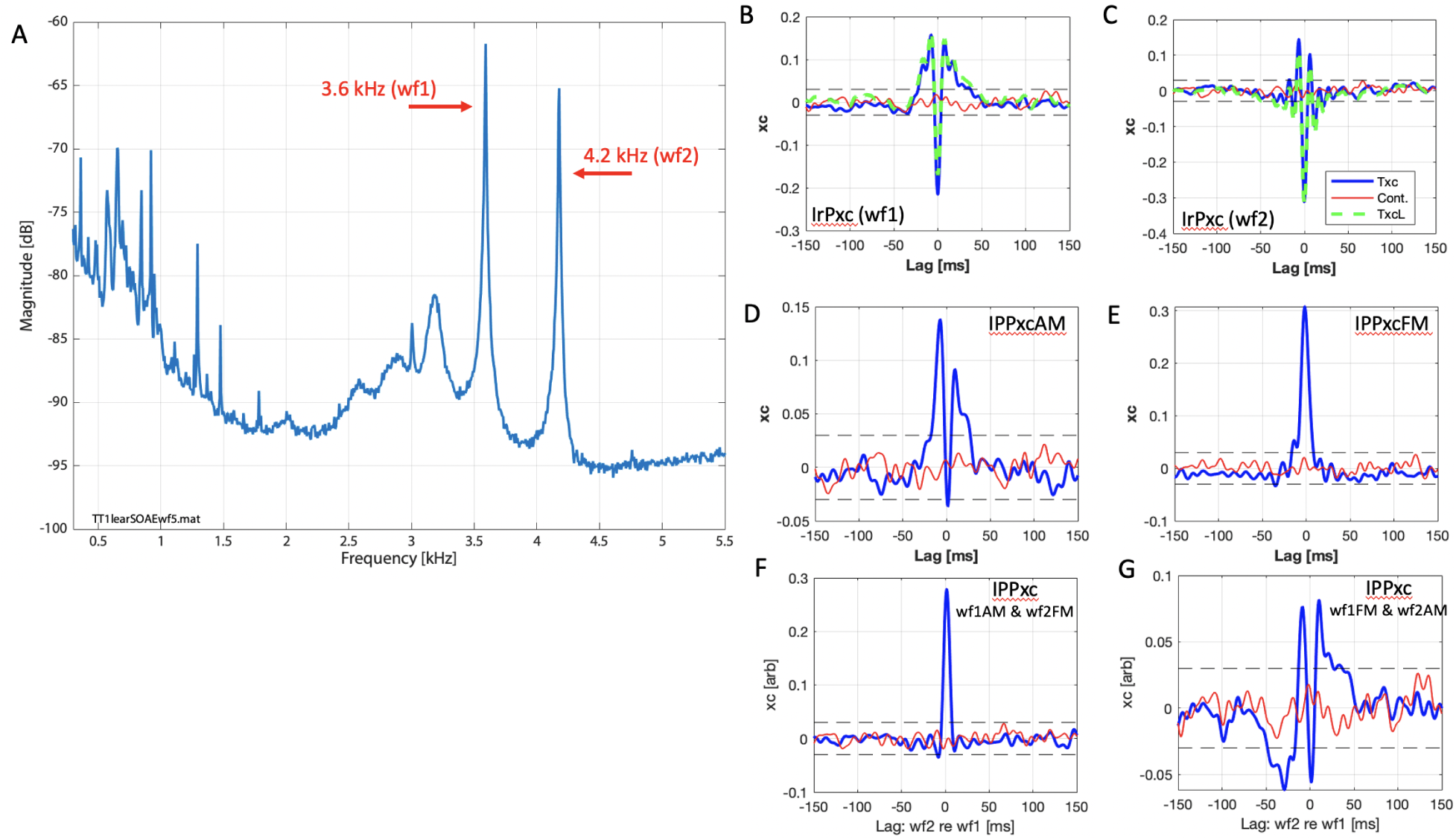
Example SOAE responses and corresponding correlative behavior for a tegu (*Salvator merianae*). Panel A shows the SOAE spectrum, with peaks at 3.575 and 4.177 kHz flagged for analysis. Both peaks were identified as bimodal. Note that a broad baseline between 2.2 and 4 kHz is also readily apparent. Panels B–G show various IrP and IPP correlations, as indicated in the inset labels.

**Supplementary Figure S20:**
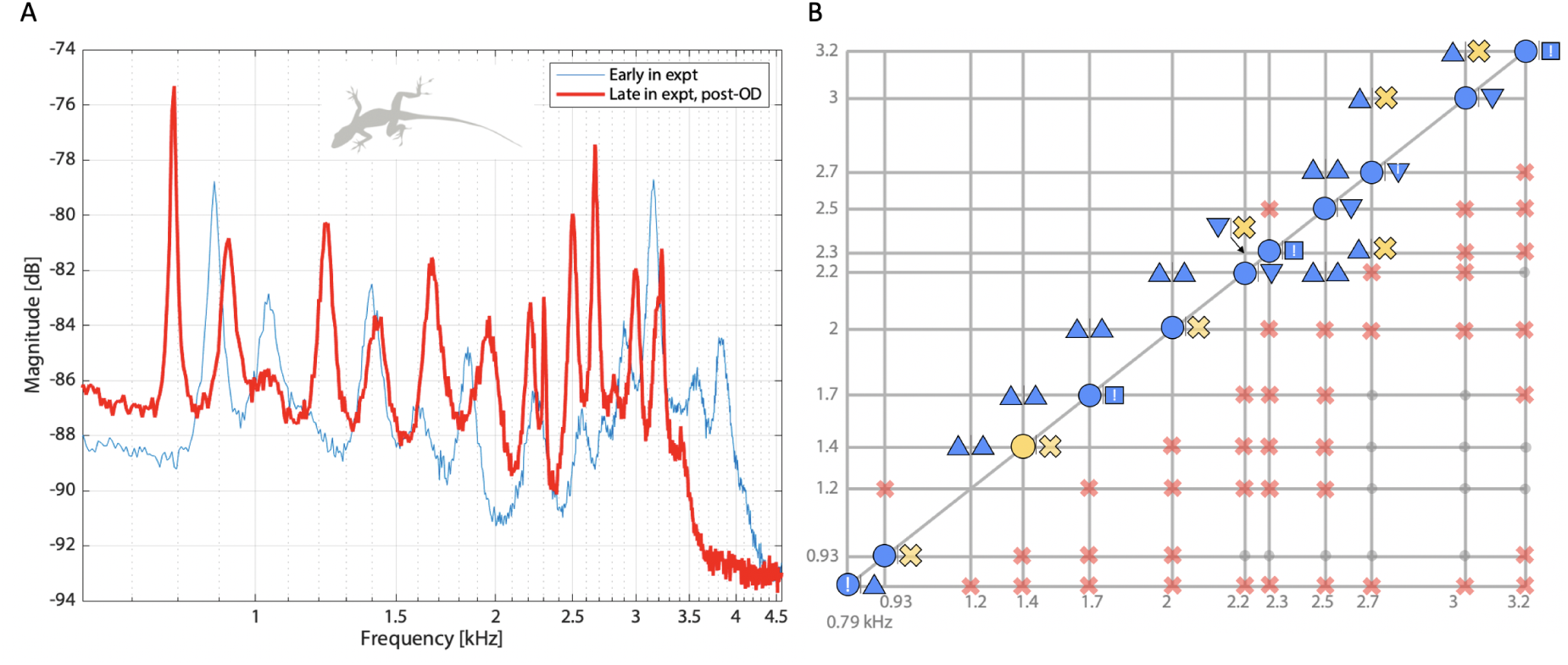
Change in SOAE spectra and correlation map upon anesthetic overdose for lizard shown in top right panel of Figs.1 and 4 (ACsb27). **A**-Spectrum (red) along pre-overdose (same as spectrum in top right panel of Fig.1) for comparison. Note the logarithmic frequency scale here. **B**-Corresponding correlation map for red curve. Same legend as Fig.4.

**Supplementary Figure S21:**
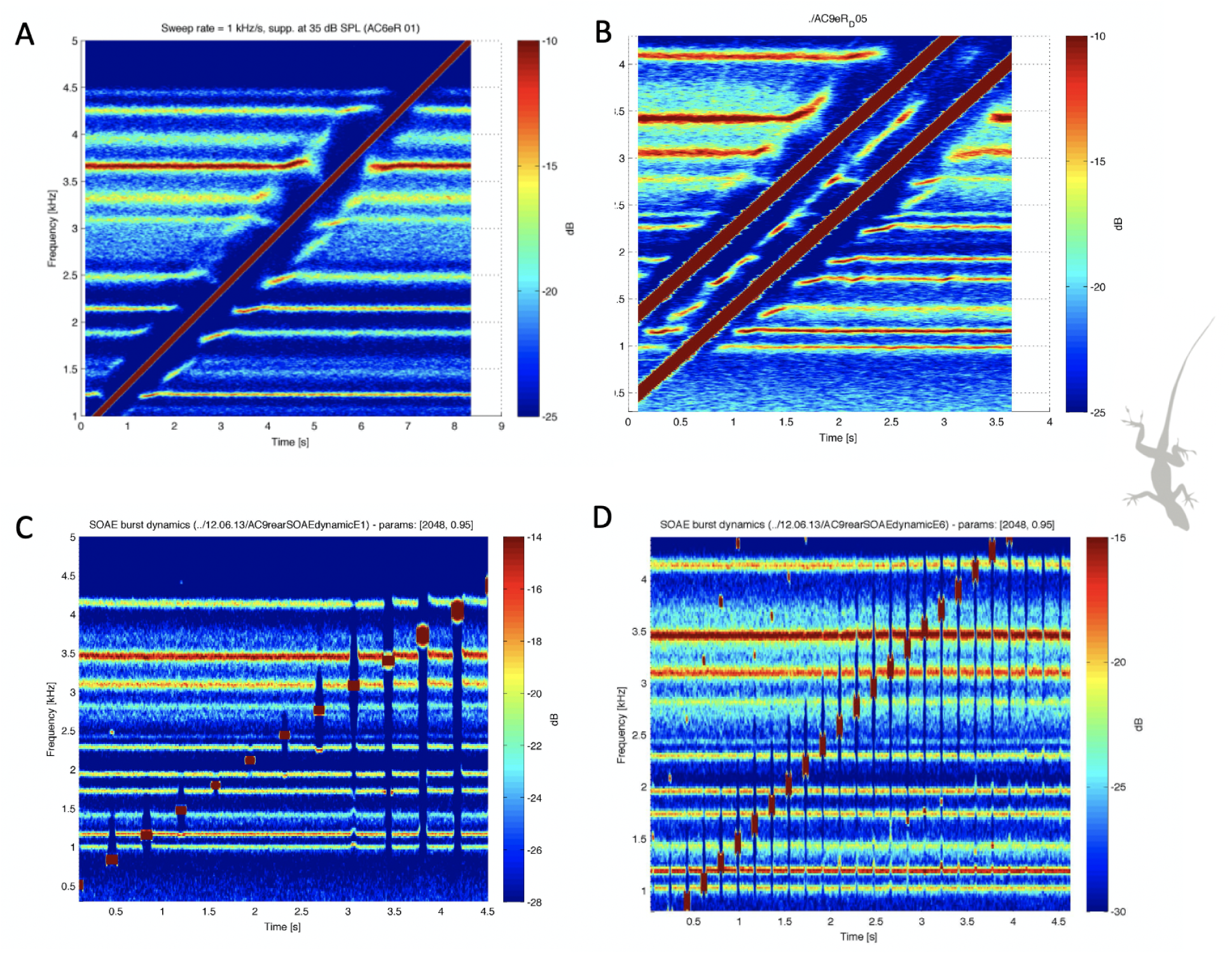
Examples of Anolis SOAE responses to a (A) moderate-level single swept tone, (B) pair of swept tones with a fixed frequency distance, (C) low-level swept tone-burst, and (D) shorter duration moderate-level swept tone-burst. See (Bergevin and Salerno, 2015) for further details.

## Footnotes

1 The term “spontaneous otoacoustic emission” (and acronym SOAE) is commonly used in a plural sense (e.g., SOAEs *≡* spectral peaks from a given ear), conflating the fact that SOAE peaks can be considered both narrow and broadband. In an attempt to clarify the biophysical underpinnings, we adopt here a convention that SOAE is inherently singular in that the term refers to the underlying processes and associated emission of sound from the inner ear. More specifically, we take the grammatical stance that SOAE *≡ the (potentially broadband) sound emitted from the inner ear*.

2 The notion of cooperativity is a general one in that it can arise at different spatial scales, such as the molecular level (e.g., collective effort of MET channels in a given bundle; Gianoli et al., 2022), or meso-cellularly (e.g., bi-directionally oriented hair cells at a given longitudinal location in the lizard; Beurg et al., 2022). More colloquially, cooperativity indicates that the ear as whole is different from the sum of the parts.

3 Possible sources of noise that could cause the observed fluctuations include noise internal to the ear (e.g., Brownian motion of hair cell bundles due to thermal fluid agitation) that could act locally and/or globally, stochastic aspects of active force generation (e.g., channel clatter, Sasmal and Grosh, 2018), and external sounds (e.g., intrinsic thermal motion of air molecules, respiration) that could affect eardrum motion (or act through bone conduction) and thereby affect the inner ear.

4 The choice of parameters for computing the FFT can affect the absolute peak height but not the overall spectral structure (see SI, specifically Fig.S1).

5 By conservative, we mean that the use of Hartigans’ dip statistic is underestimating the fraction of bimodal peaks, given limitations inherent to the method itself. This is justified by visual examination of the (filtered peak) analytic signal as a 2-D histrogram in the complex plane. For example, inspection suggests that nearly 2/3 of anole SOAE peaks are bimodal, rather than about 1/4 as indicated in Table 3.

6 The Fourier transform of a rectangular pulse is a sinc function. We note that though the (recursive exponential) filter shape employed in this study is not rectangular per se (see green curve in Fig.S4A), it is rectangular-like enough to expect this ringing behavior to first order.

7 For example, can the expanded size and more complex morphology of human and owl relative to anole be indicative of richer dynamics and interactions occurring (or not) over different timescales in those ears? Further, to what extent can different (stronger?) inter-element coupling via the TM explain the present results?

8 There is an error in Negandhi et al. (2018). There is a passage that reads “The length of the organ is 3.5 mm”. Their Fig.3 clearly shows that value should be 0.35 mm, or 350 *µ*m.

9 An analogy may be useful here. There are ubiquitous real-world examples of “global oscillators”, where the system-level spontaneous oscillation arises despite that the constituent parts themselves would not if in isolation. A laser cavity is one, to which the cochlea has been likened (Shera, 2003). Another illustrative case would be an automobile. Take a car apart and you have myriad passive components. However, put them together in the correct fashion and something entirely else emerges. Even an idle car will intrinsically vibrate, which is readily perceptible (e.g. put your hand on the hood). This arises effectively due to the engine being designed to “oscillate” and rotationally drive the crankshaft. While not functionally relevant themselves, these vibrations arise as an epiphenomenon tied to some other purpose (i.e. making a car move).

10 Note that a limit-cycle oscillator, such as a van der Pol system, subject to strong positive damping and/or a constant force pushing it way from the region where a negative damping occurs, may not self-oscillate. Thus, there are myriad scenarios where a limit-cycle oscillator is quiescent. Stochastic forces complicate things further [e.g. Sheth et al. (2018)].

11 Waves may be various types, such as traveling versus standing waves. Further, waves could even be diffusive in nature (Mandelis, 2000). Physiologically verifying which, if any, of these wave types are present should greatly inform how precisely energy generated in one location affects generation sites at spatially removed areas.

